# Spontaneous Cell Fusions as a Mechanism of Parasexual Recombination in Tumor Cell Populations

**DOI:** 10.1101/2020.03.09.984419

**Authors:** Daria Miroshnychenko, Etienne Baratchart, Meghan C. Ferrall-Fairbanks, Robert Vander Velde, Mark A Laurie, Marilyn M. Bui, Aik Choon Tan, Philipp M. Altrock, David Basanta, Andriy Marusyk

## Abstract

Initiation and progression of cancers reflect the underlying process of somatic evolution, which follows a Darwinian logic, *i.e*., diversification of heritable phenotypes provides a substrate for natural selection, resulting in the outgrowth of the most fit subpopulations. Although somatic evolution can tap into multiple sources of diversification, it is assumed to lack access to (para)sexual recombination – a key diversification mechanism throughout all strata of life. Based on observations of spontaneous fusions involving cancer cells, reported genetic instability of polypoid cells, and precedence of fusion-mediated parasexual recombination in fungi, we asked whether cell fusions could serve as a source of parasexual recombination in cancer cell populations. Using differentially labelled tumor cells, we found evidence of low-frequency, spontaneous cell fusions between carcinoma cells in multiple cell line models of breast cancer both *in vitro* and *in vivo*. While some hybrids remained polyploid, many displayed partial ploidy reduction, generating diverse progeny with heterogeneous inheritance of parental alleles, indicative of partial recombination. Hybrid cells also displayed elevated levels of phenotypic plasticity, which may further amplify the impact of cell fusions on the diversification of phenotypic traits. Using mathematical modeling, we demonstrated that the observed rates of spontaneous somatic cell fusions may enable populations of tumor cells to amplify clonal heterogeneity, thus facilitating the exploration of larger areas of the adaptive landscape, relative to strictly asexual populations, which may substantially accelerate a tumor’s ability to adapt to new selective pressures.

## INTRODUCTION

Cancer is the direct result of somatic clonal evolution, which follows Darwinian principles: diversification of heritable phenotypes provides a substrate, upon which natural selection can act, leading to the preferential outgrowth of phenotypes with higher fitness in the specific environment^1,2^. Thus, the ability to generate new heritable diversity is required for the evolvability of populations of tumor cells, both during tumor progression and in response to therapies. Evolving tumors have access to several powerful diversification mechanisms that are considered the enabling characteristics within the hallmarks of cancer framework^3^: genomic instability, elevated mutation rates, and deregulation of epigenetic mechanisms that control gene expression. At the same time, cancer cells are generally assumed to lack a key evolutionarily-conserved source of diversification—sexual or parasexual (exchange of genetic material without meiosis) recombination. Within genetically diverse populations, (para)sexual recombination can dramatically amplify diversity and generate new mutational combinations (thus enabling new epistatic interactions), while unlinking advantageous mutations from disadvantageous ones, hence supporting population fitness and accelerating evolutionary adaptation^4–6^.

Populations of tumor cells are assumed to be strictly asexual, i.e., all novel genetic and epigenetic solutions “discovered” by tumor cells are thought to be strictly clonal, inheritable only by the direct progeny of (epi)-mutated cells. However, occurrences of spontaneous cell fusions involving tumor cells have been documented both *in vitro* and *in vivo^7–9^*. Given the previously reported impact of genome duplication on increased genomic instability^10,11^, evidence of ploidy reduction in the progeny of experimentally induced hybrid cells^12,13^, reported genetic recombination in asexual ploidy cycle of cancer cells^14^, and existence of parasexual life cycles involving fusion-mediated recombination in fungi, such as pathogenic yeast C. albicans^15^, we decided to examine whether spontaneous cell fusion could serve as a source of diversification in tumor cell populations. We found that, while relatively infrequent, spontaneous cell fusions can be detected in a wide range of breast cancer cell lines both *in vitro* and *in vivo*. A subset of these hybrid cells are clonogenically viable. Whereas cell fusion simply combines two genomes, some of the hybrids undergo ploidy reduction that is accompanied by genome recombination, which generates new sub-clonal diversity. Our *in silico* modeling suggests that this fusion-mediated recombination could augment the evolvability of tumor cell populations even when spatial limitations are considered. Thus, our studies suggest that spontaneous cell fusions may provide populations of tumor cells with a mechanism for parasexual recombination and make them capable of exploring combination of mutations from different clonal lineages, thus accelerating diversification and enhancing evolvability.

## RESULTS

In the course of multiple experimental studies involving *in vitro* co-cultures of tumor cells that carry different fluorescent protein labels, we occasionally noticed double-positive cells upon fluorescence microscopy analyses (**Fig 1a, S1a)**. Similarly, we observed double-positive cells in co-cultures of tumor cells and cancer-associated fibroblasts (CAFs) (**Fig. S1b**). Examination of time-lapse microscopy images revealed that these double-positive cells can originate from spontaneous cell fusions **(Fig. S2a, b, Videos S1-5**). The phenomenon was not limited to *in vitro* cultures. Confocal microscopy examination of experimental xenograft tumors also revealed the occasional presence of cells expressing both fluorescent labels (**Fig. 1b**).

**Fig.1.**
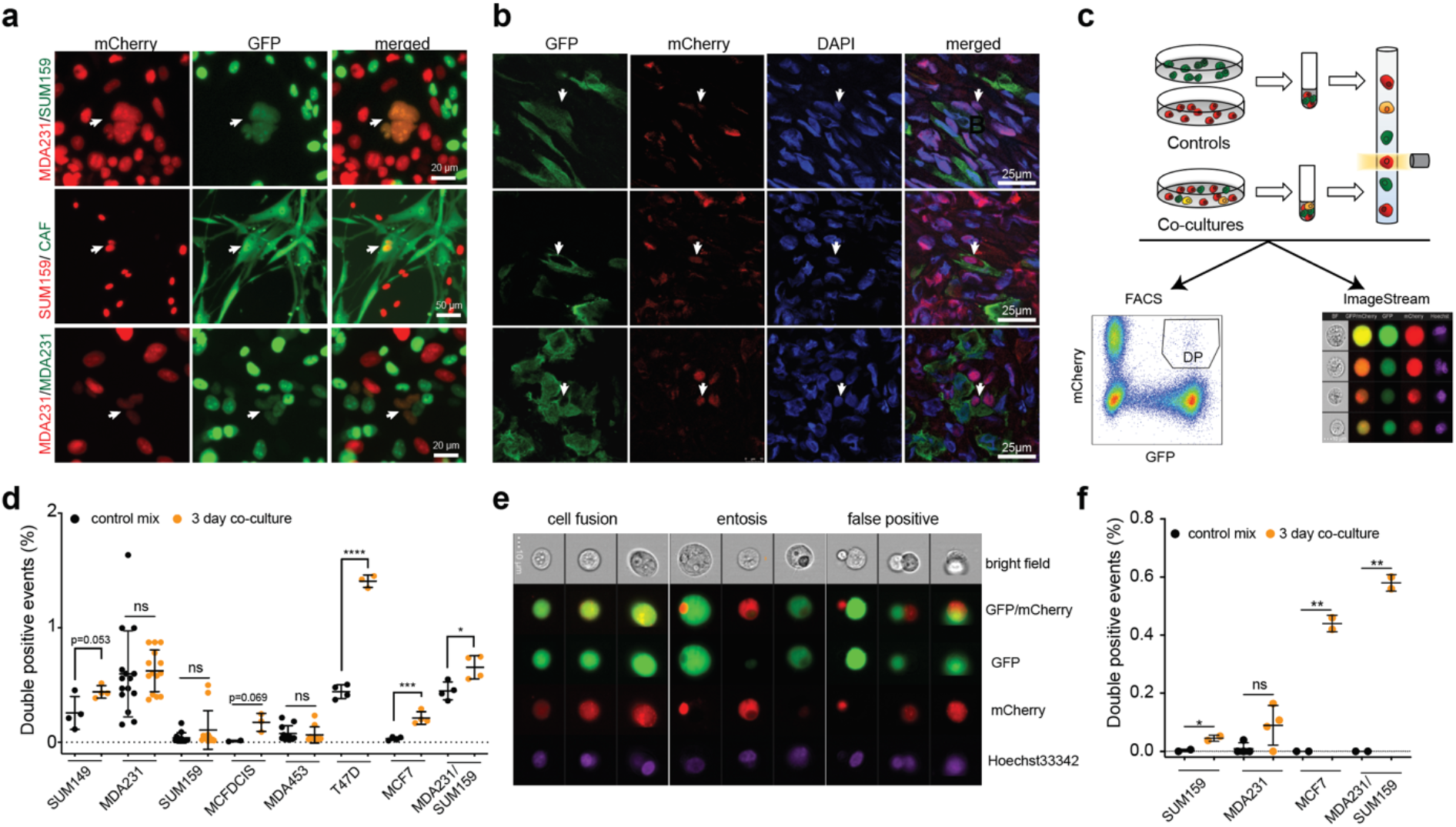
Detection of spontaneous cell fusions *in vitro* and *in vivo*. **a.** Live fluorescence microscopy images of co-cultures of the indicated differentially labelled cells. Arrowheads indicate cells co-expressing both fluorescent labels. **b.** Confocal immunofluorescent images from a xenograft tumor, initiated with a 50/50 mix of GFP and mCherry-labelled MDA-MB-231 cells. Arrowheads indicate cells co-expressing GFP and mCherry. **c.** Experiment schemata for flow cytometry (conventional and ImageStream) studies. DP in a representative flow cytometry histogram indicates double-positive (GFP and mCherry) populations. **d.** Quantification of FACS-detected frequencies of DP events of *in vitro* cell fusions of the indicated homotypic and heterotypic mixes. Each dot represents a measurement from an independent biological replicate. **e.** Representative images from ImageStream analyses of co-cultures of differentially labelled MCF7 cells. Hoechst33342 was used as a nuclear stain. **f**. Quantification of visually validated DP events from the ImageStream data. *, **, ***, *** indicate p values below 0.05, 0.01, 0.001, and 0.0001, respectively, for two-tailed unpaired t-test.

Given the possibility that spontaneous cell fusions between genetically distinct cells might provide evolving populations of tumor cells with a new source of genetic diversification, we decided to systematically investigate this phenomenon. To this end, we labelled panels of breast cancer cell lines and primary breast CAF isolates with lentiviral vectors expressing GFP and mCherry reporters and co-expressing antibiotic resistance markers blasticidin and puromycin, respectively. Differentially labelled cells of the same (homotypic) or distinct (heterotypic) cell lines were plated at a 1:1 ratio and, after co-culturing for 3 days, subjected to flow cytometry analysis (**Fig. 1c**). Compared to the separately cultured controls, harvested and admixed no more than 30 min prior to the analysis, all of the heterotypic co-cultures and five out of seven examined homotypic cultures exhibited higher proportions of events in the double-positive gate (two of the homotypic cultures reached statistical significance) (**Fig. 1d, Fig. S2c**).

The significantly higher proportion of double-positive events detected by flow analysis than that by microscopy examination, substantial within group variability, and the detection of double-positive events in some of the negative control samples indicated significant rates of false positives. Therefore, we set to validate the flow cytometry findings using ImageStream, an imaging-based platform that combines the high processivity of flow cytometry analysis with the ability to evaluate recorded images of each event^16^. Indeed, examination of the images of double-positive gate events (gating logic provided in **Fig.S3a**) revealed significant rates of false positives reflecting cell doublets (**Fig 1e**). Some of the double-positive events were cell-within-cell structures, indicating entosis^17^ or engulfment of cell fragments (**Fig 1e**). Still, a substantial fraction (~20%) of double-positive events were unambiguous mono- or bi-nucleated single cells with clear, overlapping red and green fluorescent signals, indicative of *bona fide* cell fusions (**Fig 1e, f, S3b)**. Direct comparison of flow cytometry and ImageStream analysis of the same sample revealed that true positives represented ~30% of the double-positive events detected by flow analysis (**Fig. S3c**). Consistent with the expected increase in cell size resulting from the fusion of two cells, the double-positive cells were significantly larger than cells expressing a single fluorescent marker (**Fig. S3d**). In summary, these results suggest that, while spontaneous cell fusions between cancer cells are relatively infrequent, they occur in a wide range of experimental models.

Next, we asked whether hybrid cells, formed by spontaneous somatic cell fusions, are capable of clonogenic proliferation. To this end, we co-cultured the differentially labelled cells for 3 days and then subjected them to the dual antibiotic selection (**Fig. 2a**). After two weeks of selection, which was sufficient to eliminate cells in the single-labelled negative controls, all of the examined breast cancer cell lines invariably contained viable, proliferating cells expressing both GFP and mCherry fluorescent markers (**Fig. 2b**). Notably, the clonogenic proportions of cells with dual-antibiotic resistance was lower than the frequency of fusion events (**Fig. 1f** and **2c**), suggesting that only some of the hybrids were capable of sustained proliferation. Similarly, we were able to recover dual-antibiotic-resistant cells expressing both fluorescent markers from *ex vivo* cultures of xenograft tumors initiated by co-injection of differentially labelled cells (**Fig. 2b**). Surprisingly, despite the relatively high rates of fusion detected by flow cytometry (**Fig S2c**) and microscopy, as well as previous reports on formation of viable hybrids formed by fusions between carcinoma and stromal cells^18,19^, we were unable to recover colonies from co-cultures between multiple breast cancer cells and three distinct primary, non-immortalized CAF isolates.

**Fig.2.**
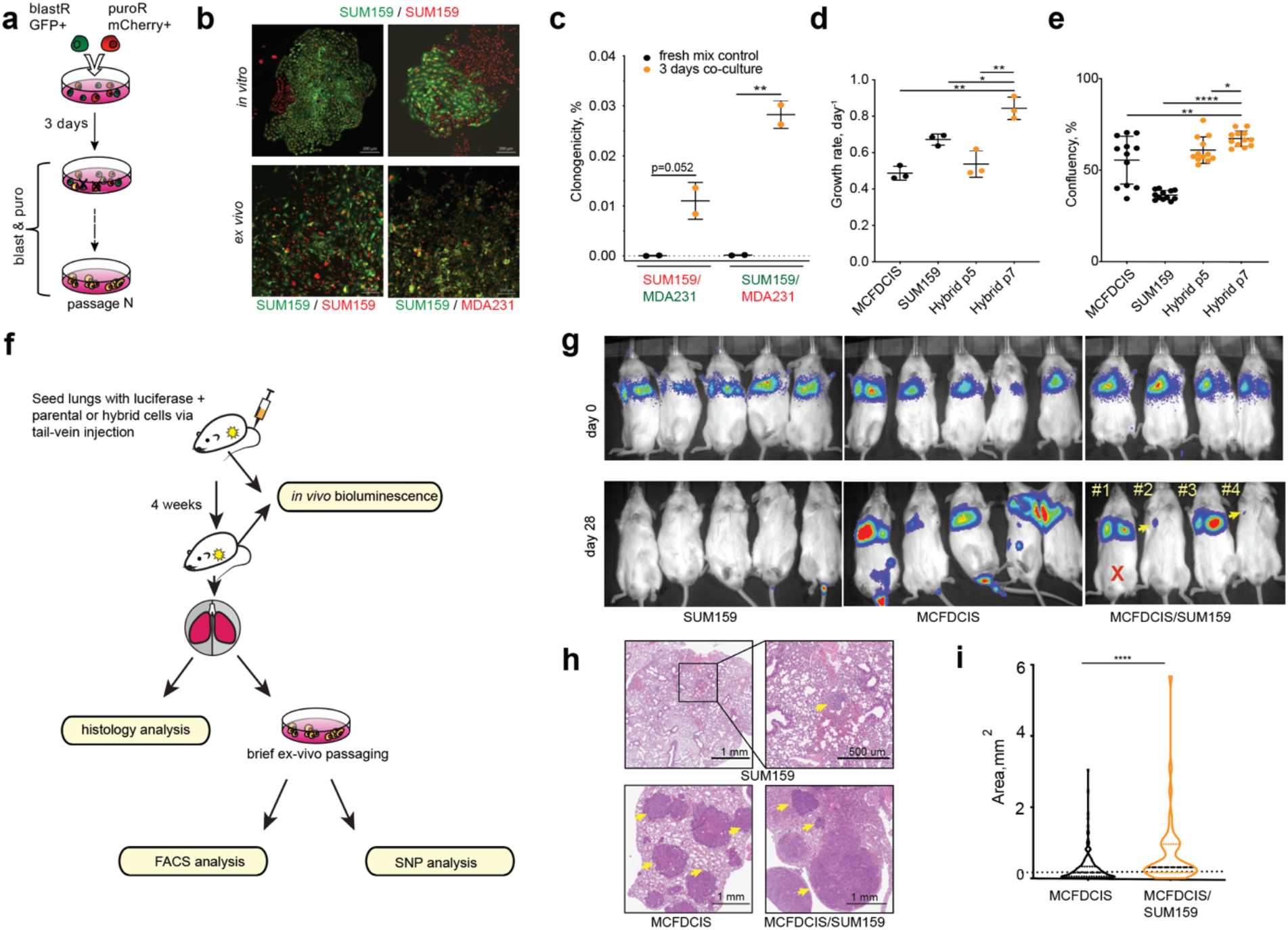
Phenotypic characterization of hybrids. **a.** Experiment schemata for the selection of hybrid cells. **b.** Representative images of live fluorescent colonies formed after selection of the indicated *in vitro* co-cultures or *ex vivo* tumors. **c.** Quantitation of frequency of fusions leading to clonogenically viable hybrid cells. **d.** Growth rates of the indicated parental cell lines and hybrids at the indicated passages. **e.** Quantification of transwell cell migration assays of the indicated cells. **f**. Experiment schemata for analyses of lung colonization. g. *In vivo* bioluminescence imaging of animals, injected with mixed SUM159PT/MCFDCIS hybrids or parental cells via tail vein injection to seed lung metastases. #1-4 indicate individual animals, used for subsequent analyses. Red X denotes a mouse that died prior to euthanasia. **h.** Representative images of H&E stains of lungs from the indicated xenograft transplants. Yellow arrows point to example tumors. Inlet showed magnified image of a micrometastasis. **i.** Violin plot of size distribution of individual macrometastatic lung tumors from the analyses of H&E stained histology slides. *, **, ***, *** indicate p values below 0.05, 0.01, 0.001 and 0.0001, respectively, for the two-tailed unpaired t-test (c, d& e) and Mann-Whitney U test (h).

Next, given the prior reports of fusion-mediated increase in invasive and metastatic potential^19,20^, we asked whether hybrids formed by spontaneous somatic fusions between cancer cells differ in their proliferative and invasive potential from those of parental cells. At early passages during the post-antibiotic-selection phase, hybrids displayed lower net proliferation rates than did the parental cell lines (**Fig. 2d, Fig. S4a**). However, at later passages, most of the examined hybrids matched and, in some cases, exceeded the proliferation rates of the fastest growing parent. This observation of an increase in proliferation rates with passaging is consistent with the elimination of viable but non-proliferative hybrids along with the selection of variants with higher proliferative abilities (**Fig. 2d, Fig. S4a**). Transmembrane invasion assays revealed that most hybrids displayed invasive rates equal to or exceeding rates of the more invasive fusion parents (**Fig. 2e, Fig. S4b, c**).

Next, we assessed the impact of somatic cell fusions on metastatic colonization potential. To this end, we compared lung colonization potential between a cell line with a relatively weak lung colonization potential (SUM159PT) and a cell line with a strong potential (MCF10DCIS), along with their hybrids using the tail vein injection assay (**Fig. 2f)**. Despite identical initial lung seeding efficiencies, mice injected with SUM159PT cells lost luminescent signal from the lungs (**Fig. 2g)**, although post-mortem histological examination revealed the presence of multiple micro-metastatic nodules, suggesting a microenvironmental growth bottleneck rather than inability to seed lungs *per se* (**Fig. 2h)**. In contrast, luminescent signals in all four mice injected with MCF10DCIS cells and in two out of four mice injected with the hybrid cells increased over time, while the other two hybrid cell recipients displayed reduced but detectable luminescent signal (**Fig. 2g)**. Histological examination revealed that lungs of all of the MCF10DCIS recipient and hybrid cell recipient mice contained macroscopic tumors in the lungs. Surprisingly, despite weaker luminescent signals in two out of three analyzed animals, histological examination revealed larger tumors than those in mice transplanted with MCF10DCIS cells (**Fig 2h, i, S4 d**), likely reflecting the loss of luciferase gene expression in some of the hybrids. One of the mice with strong luminescent signal (#1 in **Fig. 2f)** had died prior to euthanasia; necropsy analysis revealed massive tumors in the lungs, but due to poor tissue quality, this animal was excluded from the analysis. Notably, flow cytometry analysis of lungs recovered from the recipients of hybrid cells, revealed that the majority of fluorescent cells expressed both GFP and mCherry, including lungs of the animals that displayed a reduction in luminescent signal (#2 and #4, **Fig. 2g** and **S5a**). In summary, consistent with previously reported observations, these data suggest that that spontaneous cell fusions between neoplastic cells can generate cells with more aggressive oncogenic properties.

In the absence of the TP53 dependent checkpoint function, which is commonly disrupted in cancer cells, polyploidy is known to be associated with increased genomic instability^11^. Consistently, genomic instability^21^ and ploidy reduction^12,13^ were reported in experimentally-induced somatic hybrids. Therefore, we decided to examine whether spontaneously formed hybrid cells can maintain stable genomes over time. As expected, at early (1-4) passages, counted after the complete elimination of singe-antibiotic resistant control cells, all of the examined hybrid cell lines displayed elevated DNA content, consistent with the combined genomes of two parents (**Fig. 3b, Fig. S6**). However, the average ploidy of three out of seven examined hybrids was evidently reduced with additional passaging (passages 4-10), while the average ploidy of the remaining four hybrids remained seemingly unchanged (**Fig S6**). Given that genomic instability of tumor cells can be enhanced by genome doubling^22^, that fusion-mediated recombination and the stochastic loss of parental DNA accompanying ploidy reduction can serve as the mechanism for parasexual recombination in the pathogenic yeast species C. albicans^15,23^, and that cycles of somatic cell fusions followed by genetic recombination and ploidy reduction have been described to operate in normal hepatocytes^24^ and hematopoietic cells^25^, we decided to compare the genomes of single-cell-derived subclones of somatic hybrids (derivation schemata shown in **Fig. 3a**). Consistent with the maintenance of polyploid genomes in mixed populations, all of the examined sub-clonal derivatives of SUM159PT/MCF10DCIS and MDA-MB-231/MCF10DCIS hybrids retained elevated DNA content (**Fig S6**). In contrast, the genomes of individual subclones from the hybrids with reduced ploidy (MDA-MB-231/Hs578T and MDA-MB-231/SUM159PT) displayed substantial variation in DNA content (**Fig 3b, Fig. S6),** suggestive of genomic diversification.

**Fig.3.**
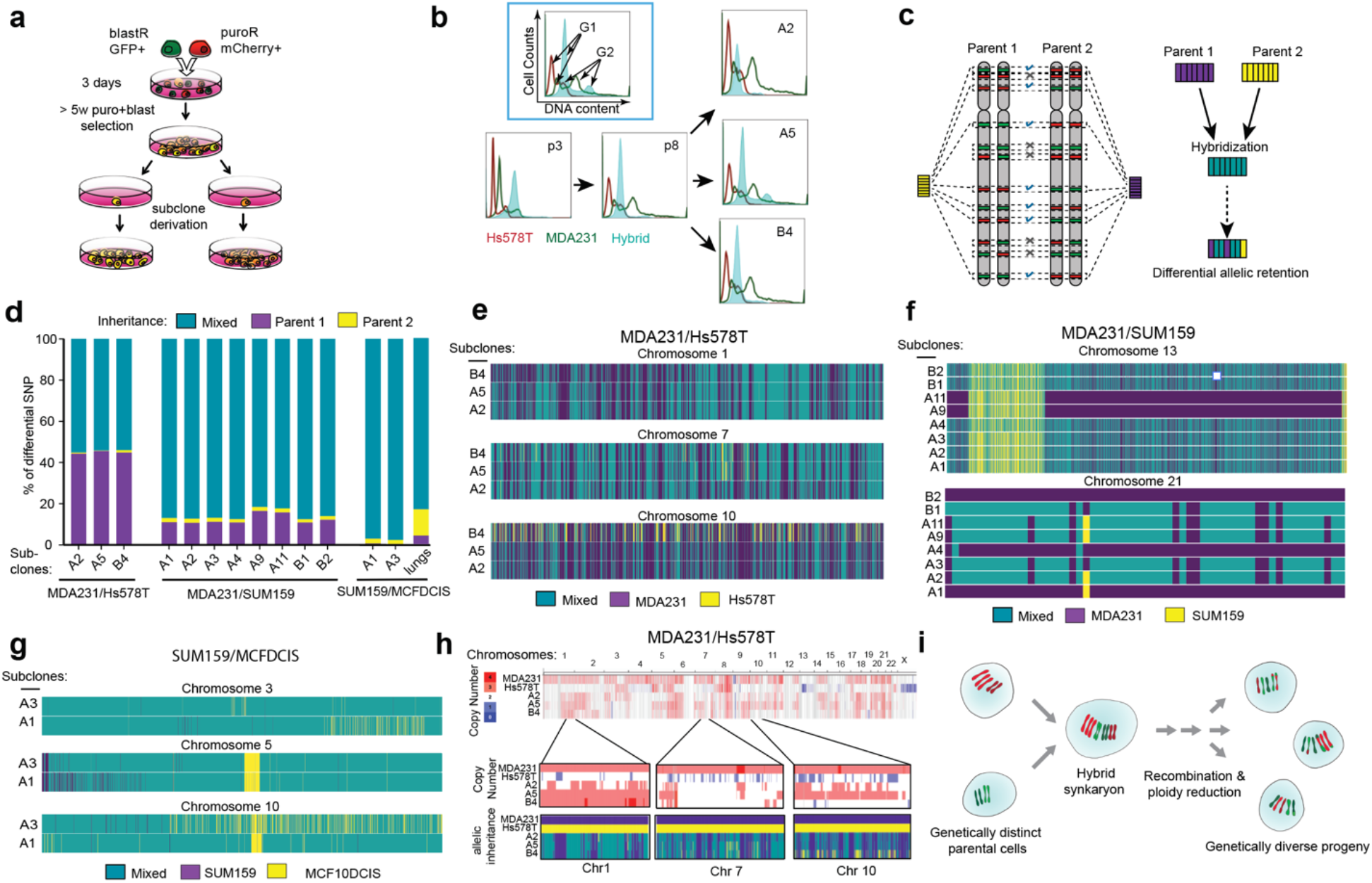
Fusion-mediated genetic diversification. **a.** Experiment schemata for the derivation of hybrid subclones. **b.** DNA content analysis of parental cell lines (in this example, HS578T and MDA-MB-231) and hybrids. PI based DNA content profiles of the hybrids are superimposed with profiles of the parent cells. Inlet describes the axes and indicates positions of the G1 and G2 peaks. A2, A5, B4 denote distinct hybrid subclones; p3 and p8 refer to the passage number of the mixed hybrid population. Shift of the DNA content profile to the left with extended passages indicates reduction of average cell ploidy in the hybrids. Differences in the profiles of individual hybrid subclones indicate diversification in DNA content. **c**. Experiment schemata of the pipeline for the analysis and visualization of SNP inheritance. **d**. Summary of analyses of inheritance of cell line-specific SNP alleles across the indicated hybrid subclones. “Lungs” refers to the mixed population, isolated from colonized lungs of mouse #3 from Figs. 2g & S5. **e-g.** Inheritance of cell line-specific alleles mapped to specific chromosomes in the hybrid subclones of the indicated hybrids. Columns represent individual alleles, with color code indicating mixed or parent specific inheritance. Maps for all of the individual chromosomes are provided in Figs. S7-9. **h**. Analyses of copy numbers for the cell line-specific alleles in parental cell lines and hybrid subclones. Selected zoom-ins of illustrative chromosomal regions depict correspondence between copy numbers and inheritance of cell line-specific SNPs. i. Schema of a proposed fusion-mediated diversification.

In order to gain deeper insights into the impact of hybridization on genomic diversification, we compared patterns of inheritance of cell-line-specific alleles. To this end, we characterized single nucleotide polymorphism (SNP) profiles of the genomically unstable MDA-MB-231/Hs578T hybrids and the parental cell lines using the Affymetrix CytoScan SNP platform. Focusing on homozygotic cell line-specific SNPs that differentiate the parental cells, we characterized their inheritance in individual single cell-derived hybrid subclones, isolated from mixed populations of hybrid cells (**Fig. 3c**). Genome-wide analysis of allelic inheritance revealed that for the majority of the SNPs that discriminate the parental cell lines, the subclones showed mixed inheritance. However, in 44.9 +/− 0.4% only MDA-MB-231 specific alleles were detected; in a smaller fraction of the analyzed loci (0.62 +/− 0.26%), only Hs578T unique alleles could be detected (**Fig. 3d, Table S1**). Importantly, differential inheritance was not confined to whole chromosomes or large chromosomal regions. Instead, we observed a significant mosaicism in the SNP inheritance within individual chromosomes, with substantial variability in the degree of mosaicism between individual chromosomes (selected examples are shown in **Fig. 3e**, with a complete set of chromosomes shown in **Fig. S7**). Individual subclones displayed distinct patterns of inheritance of parent-specific alleles. Although a high degree of genomic rearrangements within parental cell lines complicates the analyses, mosaic inheritance of parental SNPs across individual chromosomes and substantial variability between distinct subclones strongly suggest that ploidy reduction has been accompanied by partial genomic recombination.

We observed similar patterns of mosaic loss of parent-specific alleles, as well as variability in patterns of inheritance between distinct subclones in MDA-MB-231/SUM159PT hybrids, analyzed with the Illumina CytoSPN-12 platform (**Fig. 3d, f, Fig. S8**). Notably, analysis of the SUM159PT/MCF10DCIS hybrids, where both fusion parents have relatively stable, near-diploid genomes and the hybrid populations do not show obvious signs of ploidy reduction (**Fig. S6**), also revealed the loss of some of the parent-specific alleles, as well as divergence in allelic inheritance between the two distinct subclones that we have analyzed, although to a lower extent than that in the MDA-MB-231/Hs578T and MDA-MB-231/SUM159PT hybrids (**Fig. 3d, g, Fig. S9)**. Interestingly, analysis of hybrid cells, recovered from tumor-bearing lungs of a mouse (#3 shown in **Fig. 2g**) injected with pooled SUM159/MCF10DCIS hybrids revealed distinct and more extensive patterns of inheritance of parental SNPs (**Fig. S5b**), potentially reflecting the impact of distinct selective pressures experienced by cells *in vivo*.

While we observed a substantial interclonal variability in patterns of mosaic SNP inheritance, a large fraction of SNPs displayed identical patterns of inheritance in distinct subclones. Given that the numbers of distinct SNP alleles can vary between highly aneuploid genomes of different cancer cell lines, this unequal contribution could be the most parsimonious explanation for the observed similarities in the patterns of SNP inheritance. Should this be the case, more numerous alleles would be more likely to be retained in hybrids under stochastic ploidy reduction. Additionally, this numeric inequality could lead to detection issues, where numerically superior alleles from one parent could potentially mask the signal from less numerous alleles from another parent. Differences in the numbers of cell line-specific alleles inherited from different parental cell lines might also explain the apparent dominance of the MDA-MB-231 cell lines in the two hybrids that we have analyzed by SNP arrays. To address whether distinct copy numbers of differential SNP alleles could explain the observed inheritance patterns, and to examine the variegation in allelic copy numbers between distinct subclones, we contrasted copy number data with allelic inheritance data across several chromosomal regions within MDA-MB-231/Hs578T hybrids. Dominance of MDA-MB-231 – exclusive and mixed allelic inheritance in MDA-MB-231/Hs578T hybrids was generally consistent with higher copy numbers of differential SNPs inherited from MDA-MB-231 cells (**Fig. 3h**). Importantly, we observed significant variability in allelic copy numbers between distinct subclones of the hybrids, indicating additional diversification. Some of this variability was consistent with differences in allelic inheritance. For example, lower copy numbers from the region of chromosome 10 in subclone B4 were linked with distinct patterns and higher proportion of Hs578T alleles (**Fig. 3h**). On the other hand, the majority of similarities and dissimilarities in patterns of allelic inheritance within different hybrid subclones could not be fully explained by differences in allelic copy numbers between the genomes of different parental cell lines, suggesting the contribution of additional factors. Analysis of the more stable SUM159/MCFDCIS hybrids revealed similar clonal variegation in allelic copy numbers and patterns of allelic inheritance that could be partially explained by unequal copy numbers of SNP alleles in the parental cells (**Fig. S10a**).

The above analyses were performed on subclones derived from the same pool of hybrids. Therefore, the distinct subclones could have been the progeny of same original fusion. To test whether similar patterns of allelic inheritance could be observed in independently derived hybrids, we derived three new hybrid subclones from each of the two independent mixed populations of MDA-MB-231/SUM159 hybrids. Despite the substantial variegation in allelic inheritance and copy numbers within distinct subclones, most of the conserved patterns of allelic inheritance were shared between subclones, derived from distinct hybrid parents (**Fig. S10b, S11**). These recurrent patterns were generally consistent with the unequal contribution of SNP copy numbers from parental genomes. However, similar to the observations in the MDA-MB-231/Hs578T hybrids, the unequal contribution of allelic copy numbers could not fully explain the observed patterns (such as mosaic patterns of mixed and SUM159-specific inheritance within chromosome 13 in **Fig. S10b**), suggesting the contribution of additional mechanisms.

In addition to genetic diversification, heterogeneity in biologically and clinically important phenotypes of cancer cells is shaped by epigenetic mechanisms^26^. Theoretical studies have suggested that cell fusions between genetically identical but phenotypically distinct cells could create significant diversity due to the resultant collision of gene expression networks^27^. Therefore, we decided to examine the impact of somatic fusions on phenotypic diversification. To this end, we performed single-cell expression profiling (10x Genomics platform) to examine the phenotypes of MDA-MB-231, SUM159PT cells and MDA-MB-231/SUM159PT hybrids at early (2) and extended (10) passages under dual-antibiotic selection (**Fig. 4a**). UMAP clustering^28^ of single-cell expression profiles revealed that the phenotypes of hybrid cells were distinct from that of both parents. Interestingly, we observed a substantial shift in the phenotypes of hybrids at the later passage, which is consistent with the selection of a fit subpopulation of hybrid cells and additional diversification (**Fig. 4b, c**).

**Fig 4.**
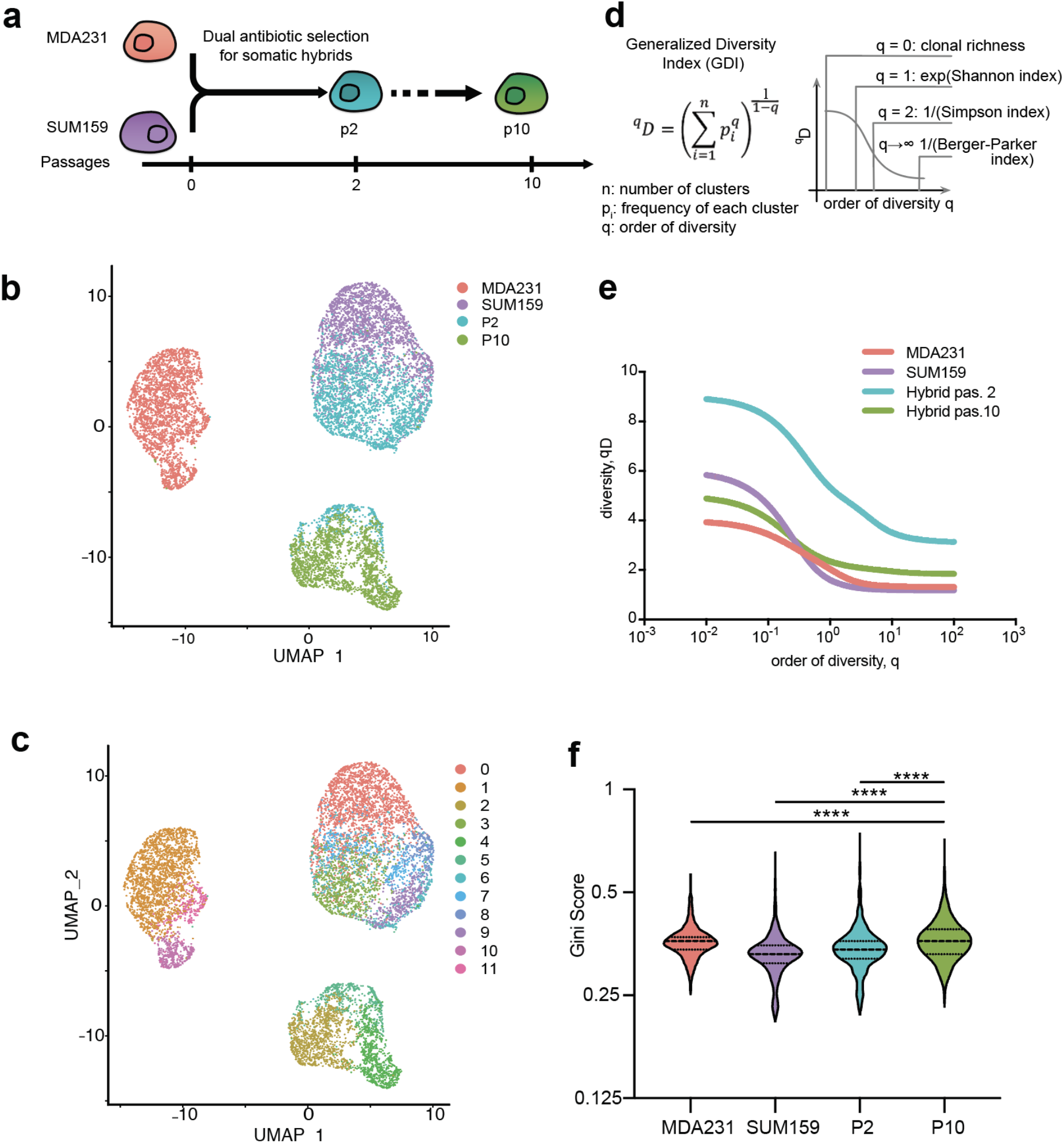
Fusion-mediated phenotypic diversification. **a.** Experimental schemata for samples used for scRNAseq analyses. **b.** UMAP distribution of cell phenotypes of parental cells and hybrids at the indicated passages from single cell expression analysis data. **c**. UMAP-defined distinct phenotypic clusters, used in GDI analyses. **d**. Formula for calculation of a generalized diversity index (GDI), and mappings to common diversity indexes that are special cases. **e**. GDI analysis of phenotypic diversity of parental and hybrid cells. **f.** Comparison of Gini scores across all of the genes with expression value >1 read in all four cell lines, between the indicated cells. Dashed lines represent medians and dotted lines represent quartiles. **** denotes p ≤ 0.0001 in a Wilcoxon signed-rank test.

To quantify phenotypic diversity within parental cell types and hybrids from the single-cell profiling data, we decided to use a general diversity index (GDI) (**Fig. 4d**)^29^. GDI enables characterization of diversity across a spectrum of orders of diversity^30^, ranging from clonal richness (low order of diversity reveals the number of distinct subpopulations) to classic measures of species diversity, such as Shannon or Simpson indices^31^ (intermediate orders of diversity), and to high orders of diversity that are giving increased weight to the highly abundant subpopulations^29^. Considering individual UMAP defined clusters as subpopulations (“species”) we found that at the early passage, hybrids displayed higher diversity across all orders of diversity (**Fig. 4e**). However, at passage 10, diversity at low orders (“species richness”) decreased. Yet, at intermediate and high orders (“species evenness”), the diversity of late passages remained higher than in either those of the parental cells.

Our GDI analyses rely on grouping phenotypes into distinct clusters. However, these analyses might miss lower level cell-to-cell phenotypic variability. Therefore, we decided to interrogate the dispersion of transcript reads across cells using the Gini dispersion index, which captures the variability of gene expression across all of the transcriptome and has been recently applied towards the characterization of phenotypic diversification in cancer cell populations^32^. We found that the MDA-MB-231/SUM159PT hybrids displayed elevated Gini indexes compared to both parents; in contrast to GDI metrics, lower level phenotypic diversity increased at the later passage (**Fig. 4f**). These findings further support the notion that somatic hybridization can lead to phenotypic diversification.

Despite the substantial genetic and phenotypic diversification observed in hybrid cells and their progeny, spontaneous fusion events are relatively rare. Whereas it is easy to intuit the potential impact of rare events creating cells with dramatically enhanced oncogenic properties, the impact of low frequency fusion-mediated recombination events on mutational diversity within tumor cell populations is less obvious. To evaluate this impact, we used *in silico* simulations based on a birth-death branching model of tumor growth (**Fig. 5a and Mathematical Supplement**). We started by deriving the probability of clonogenic cell fusions and cell proliferation from our experimental data (**Mathematical Supplement, Table S2**). Using these estimates together with values of genetic mutation rates from the literature^33^, we compared the accumulation of diversity between the scenarios of populations of tumor cells evolving through mutations only with those evolution involving mutations and fusion-mediated recombination. We found that fusion-mediated recombination can substantially enhance the increase in clonal richness (groups of tumor cells defined by unique mutational combinations) while also increasing maximum numbers of mutations observed within a single lineage (**Fig. 5b-d**). Although the impact of fusion-mediated recombination was clearly captured by the commonly used Shannon and Simpson diversity indexes (**Fig. S12a, b**), GDI enabled a more informative evaluation of the effect, showing the highest impact at the lowest orders of diversity (**Fig. 5e, Fig. S12c**).

**Fig.5.**
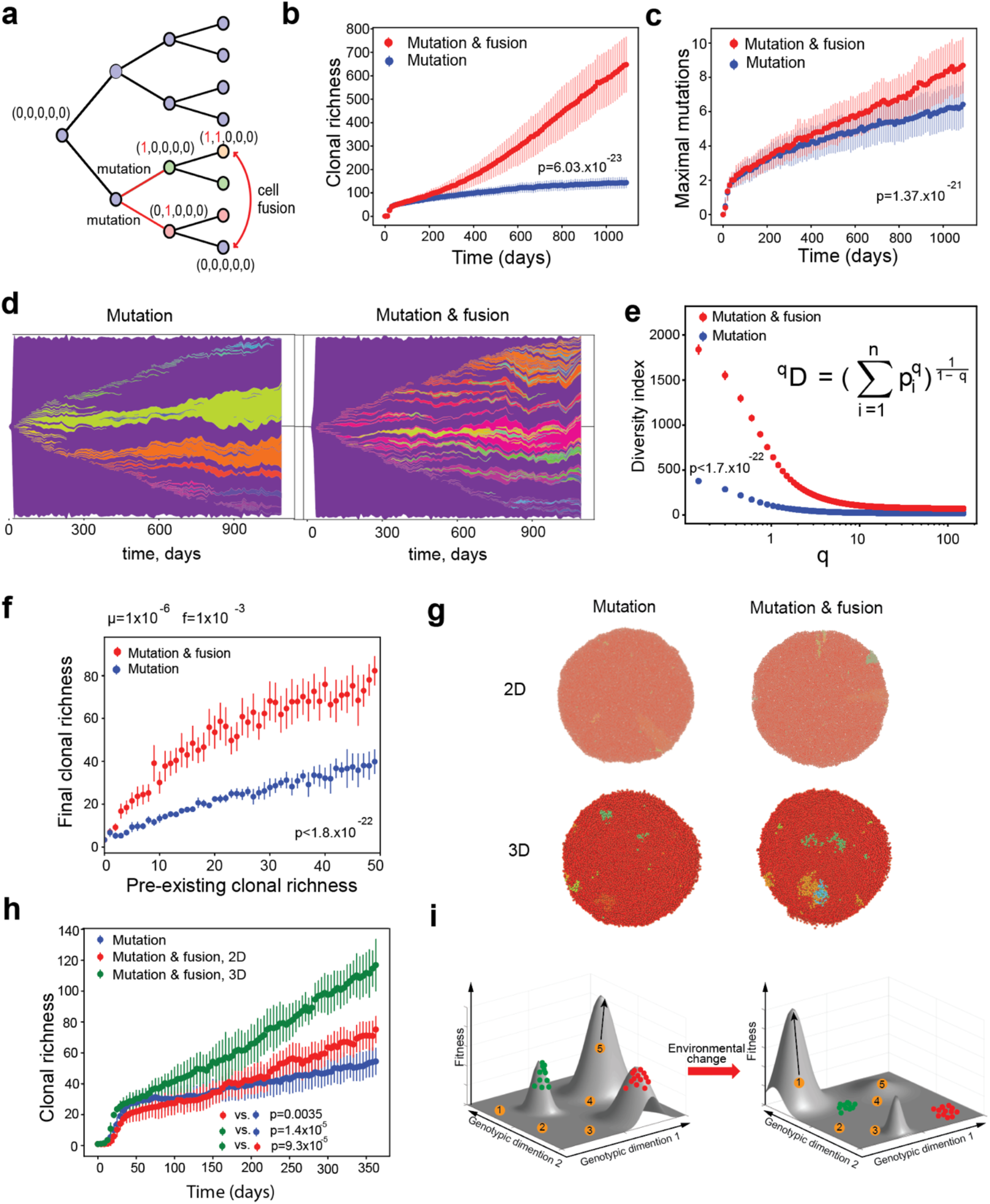
Impact of fusion-mediated recombination on genetic diversity in tumor cell populations. **a.** Schemata of the *in silico* model, using birth-death process. A binary vector represents the genotype of the cells, with (0,...,00) representing the initial genotype. Cells can stochastically acquire mutations during cell division and randomly exchange mutations during fusion-mediated recombination. Dynamics of unique mutation acquisition (**b**) and highest number of unique mutations within a single lineage (**c**) in the presence and absence of fusion-mediated recombination (fusion probability f=0.005, mutational probability μ=5×10^−5^) over 1095 days. For both (B) and (C) the solid lines represent the means and the error bars represent standard deviations of 50 simulations. **d.** Muller plot depicting the impact of fusion-mediated recombination on clonal dynamics, with f either 0 (no fusions) or 0.01, and μ=1×10^−5^. **e**. Generalized Diversity Index after 1095 days of simulation; f=0.005, μ=1×10^−4^. **f**. Impact of the initial genetic heterogeneity (clonal richness) on the ability of fusion-mediated recombination to further diversify tumor cell populations after 1095 days of in silico tumor growth at the indicated mutation and fusion rates. **g**. Visualization of the spatial distribution of subpopulations carrying unique genotypes at the end of in silico simulation of growth for 60 days with f=0.00035, μ=5×10^−3^ **h**. Number of distinct mutants over time for the spatial simulations for 365 days, f=0.00035, μ=5×10^−3^, carrying capacity 10000 cells; The solid lines represent the means over 10 seeds and the error bars are the standard deviation. All of the p values show the results of KS tests for the indicated comparisons. **i**. A model schema, illustrating the proposed impact of fusion-mediated recombination on the ability of populations of tumor cells to explore adaptive landscape. Left – under constant conditions, we assume that major genetic subclones occupy local fitness peaks and that most new variants are disadvantageous. However, some of the new mutants might “discover” a distinct fitness peak, leading to amplification through positive selection. Right – environmental change (such as initiation of therapy) changes adaptive landscape. Some of the new variants produced by fusion-mediated recombination might “discover” new fitness peaks.

This effect was observed across a biologically feasible range of fusion and mutation rates (**Fig. S13**). While larger population size and higher rates of proliferation and turnover of tumor cells predictably led to increased clonal diversity, they did not substantially modulate the impact of fusion-mediated recombination (**Fig. S14a, b**). As expected, in all cases, the impact of fusion was more pronounced at higher fusion probabilities. Less intuitively, the impact of cell fusions was also elevated by higher mutation rates. To better understand the stronger impact of fusions at higher mutation rates, we examined the ability of fusion-mediated recombination to generate new mutational variants as a function of pre-existing clonal richness, given fixed mutational and fusion probabilities. We found that higher levels of mutational heterogeneity dramatically enhanced the impact of fusion-mediated recombination (**Fig. 5f, Fig. S14c**). Therefore, the biological impact of fusion-mediated recombination should be more pronounced in tumors with higher levels of mutational clonal heterogeneity.

In our modeling thus far, we assumed a well-mixed population. However, this assumption is clearly violated in solid tumors, where spatial restrictions can have a profound impact on selective pressures and expansions of mutant subpopulations^34–37^. Thus, we asked whether fusion-mediated recombination could still have a substantial impact in spatially restricted contexts, where fusions between genetically dissimilar cells are less likely. Using spatial agent-based simulations (**Fig. S15 and Mathematical Supplement**), we compared diversification between spatially restricted populations that evolve through mutations and cell fusions versus those evolving through mutations only, in both 2D and 3D contexts (**Fig. 5g and Videos S6-9**). We found that even in spatially restricted contexts diversity was still acquired faster in the presence of fusion-mediated recombination. Notably, the impact of cell fusions was substantially higher in 3D than in 2D contexts, reflecting higher number of neighbors, which increased the probability of having a genetically distinct neighbor (**Fig. 5h**). In summary, despite the relatively low frequency of spontaneous somatic cell fusion and the impact of spatial constraints, fusion-mediated recombination can have a profound impact on somatic evolution, through the accelerated diversification of tumor cell populations and generation of rare mutational variants capable of exploring larger swathes of adaptive landscapes (**Fig. 5i**).

## DISCUSSION

Our findings reveal a surprisingly common occurrence of spontaneous cell fusions across multiple experimental cell line models of breast cancer. Spontaneous cell fusions between cancer cells and normal cells within the tumor microenvironment have been described by multiple prior studies from many groups^18,19,38–41^. Although the subject of fusions between cancer cells has received less attention, spontaneous fusions between cells have also been reported^20^. These studies have documented the impact of cell fusions on creating and enhancing malignant phenotypes, such as invasion, migration and metastatic dissemination, thus providing the basis for the argument that cell fusions can be an important contributor for cancer initiation and progression^7–9^. Our results are consistent with these prior findings. However, in addition to the previously proposed ability to generate cells with novel oncogenic properties, our results suggest the presence of a further, indirect impact with potentially profound implications. Given the evidence of genetic recombination and phenotypic diversification, resulting from spontaneous cell fusions between cancer cells, we posit that this fusion-mediated recombination and partial ploidy reduction can provide populations of tumor cells with a parasexual recombination mechanism, similar to fusion-mediated facultative parasexual recombination observed in pathogenic yeast C. albicans^15^. Using mathematical modeling we demonstrate that despite its relatively low frequency, this fusion-mediated recombination can have a significant impact on promoting evolution within cancer cell populations.

Cell fusions have been previously proposed to induce oncogenic transformation and tumor initiation through genome destabilization^13,21,41^. A large fraction of primary cancers shows evidence of whole genome doubling followed by partial genome loss^42^, although the contribution of cell fusions toward these tetraploid intermediates is unclear, as genome doubling can be also achieved by endoreduplication and aborted cell cycle^22^. Experimental studies have linked increased ploidy with elevated genomic instability, reflecting higher tolerance of chromosome aberrations^43^. Therefore, regardless of whether genome doubling is the result of cell fusions, endoreduplication or aborted cell cycle, it can lead to increased genome diversification, thus fueling cancer evolution. Indeed, whole genome doubling in primary tumors has been associated with significantly worse outcomes^43^. Nevertheless, fusions between genetically different tumor cells could have additional consequences to those arising from the duplication of the same genome or fusions between cancer and normal cells. Spontaneous fusions between genetically different tumor cells can combine mutations that have been acquired within independent evolutionary trajectories. Subsequently, the stochastic loss of some of the genetic material during ploidy reduction could create a variety of combinations of mutations from the genomes of fusion partners (**Fig. 3i**).

The extent of this additional diversification depends on the genetic divergence of parental cells, as well as on the extent of genomic recombination between the parental genomes. The latter can theoretically range from a complete lack of recombination, where ploidy reduction is achieved through losses of whole chromosomes, to an extensive blending observed in meiotic processes. Our analyses of inheritance of cell line-specific SNPs reveal mosaicism across chromosomes, which strongly indicates recombination at a level within these two extremes. Intriguingly, the extent of this mosaicism is highly variable between individual chromosomes (for example, compare chromosomes 13 and 21 in the MDA-MB-231/SUM-159PT hybrids depicted in Fig. 3f). This extensive mosaicism, confined to a subset of chromosomes, as well as the oscillating DNA copy number patterns in genomes of hybrid subclones (**Fig. 3h**) indicate a potential occurrence of chromothripsis – a single mutational event that involves the shattering a single or several chromosomes into a large number of pieces and the rejoining of these pieces in different patterns^44^, which would further increase genomic diversity following cell fusions. Thus, our results support the notion of partial recombination, with unequal impact in different areas of the genome.

Sexual and parasexual recombination, mediated by a plethora of diverse, independently evolved mechanisms, is near-universal across all strata of life, highlighting its essential importance for species evolution^45^. Of note, the frequency of clonogenic cell fusion that we have documented is similar to the frequency of parasexual recombination in yeast species with a facultative parasexual recombination life cycle^46^, supporting the notion of the potential importance of fusion-mediated recombination in somatic evolution. Indeed, our *in silico* modeling experiments suggest that the observed rates of cell fusions might significantly accelerate diversification within tumor cell populations. As genetic diversification, mediated by the putative fusion-mediated recombination works on combining and reshuffling mutational combinations from different subclonal lineages, its impact should be particularly important in tumors with higher levels of clonal mutational heterogeneity. By enabling cancer cell populations to explore epistatic interactions between mutations that are strictly confined to clonal lineages in asexual populations, fusion-mediated recombination facilitates the exploration of larger areas of adaptive landscapes, thus augmenting their evolvability.

In addition to genetic diversification (**Fig. 3**), we have observed an increase in phenotypic diversity in hybrid cells (**Fig. 4**). Although increased phenotypic variability might simply reflect genetic diversification, a recent theoretical study has suggested that cell fusions between genetically identical cells with distinct phenotypic states can trigger the destabilization of gene regulatory networks, increasing phenotypic entropy and enabling cells to reach phenotypic states distinct from those of the two parents^27^. In support of this view, cell fusions have been associated with nuclear reprogramming^47^ and increased phenotypic plasticity in multiple contexts^48^, while increased migration and invasiveness of hybrid cells (**Fig. S4**) have been linked with stemness^49^. Given the emergent importance of epigenetic mechanisms in metastatic colonization, we speculate that this increase in epigenetic plasticity, rather than genetic diversification or inheritance of invasion/metastatic properties from one of the fusion parents likely underlies increased metastatic potential of hybrid cells. Therefore, cell fusions might strongly enhance tumor cell heterogeneity and cancer progression independently of genetic diversification.

Multiple prior studies have described the occurrence of spontaneous, clonogenic cell fusions between neoplastic cells and non-neoplastic cells within the tumor microenvironment, including fibroblasts^18,38,40^, endothelial^39^ and myeloid^19,47^ cells both *in vitro* and *in vivo*. While we observed significant rates of cell fusions between breast cancer cells and CAFs (**Fig. S2c**), we failed to derive clonogenic progeny from these heterotypic hybrids. Since non-immortalized primary CAFs remain genetically normal^50^, lack of proliferation might reflect the dominant impact of the intact TP53 checkpoint to limit the proliferation of polyploid cells^11,51^. Indeed, examination of time-lapse microscopy images indicated that, in contrast to hybrids between carcinoma cells that frequently underwent cell division, hybrids involving fusion with primary CAFs either failed to proliferate or stopped proliferation after a single cell division (data not shown). While these results do not contradict the formation of proliferatively viable hybrids between cancer and non-cancer cells within the tumor microenvironment, documented in multiple prior studies^18,19,38–41^, they suggest that spontaneous somatic cell fusions between cancer cells that have lost intact checkpoint mechanisms to preserve genomic integrity are more likely to generate proliferatively viable progeny and thus more likely to contribute to diversification within cancer cell populations (**Fig. 5i**).

At this point, we cannot completely exclude the possibility that our observations of spontaneous cell fusions within *in vitro* and xenograft models might be irrelevant to primary human cancers. Nevertheless, the occurrence of viable spontaneous cell fusions has been documented *in vivo* in multiple animal models^19,38,39,41^. Cells with phenotypes consistent with hybrids between cancer cells and leukocytes have been detected in circulation in human malignancies^19^. Detecting spontaneous cell fusions in primary human cancers is notoriously difficult, as in most cases all of the neoplastic cells descend from the same clonal origin. Although giant polyploid cells consistent with spontaneous fusions can be observed in many human neoplasms^52,53^, the absence of genetic markers makes it currently impossible to discriminate cell fusions from endoreduplication or aborted cell division. Strikingly, two case reports have documented the occurrence of cancers that combine genotypes of donor and recipient genomes in patients that have received bone marrow transplantations^54,55^. These findings provide direct evidence for the potential clinical relevance of spontaneous cell fusions. Unfortunately, these cases are too rare for systematic interrogation. Still, given the abovementioned studies documenting spontaneous cell fusions in human malignancies, we postulate that our results warrant the suspension of the notion that cancers are strictly asexual. Rigorous testing of clinical relevance of spontaneous cell fusions would require the development of not only new detection approaches but also new experimental and bioinformatical pipelines to understand the degree of fusion-mediated recombination. These efforts might be well warranted, as, should fusion-mediated genetic recombination and phenotypic diversification prove to be relevant for human malignancies, they might have profound implication for evolvability of tumor cell populations, which underlies both clinical progression and the acquisition of therapy resistance.

## MATERIALS AND METHODS

### Cell lines and tissue culture conditions

Breast cancer cell lines were obtained from the following sources: MDA-MB-231, HCC1937, HS578T, T47D, MDA-MB-453 from ATCC MCF10DCIS.com from Dr. F. Miller (Karmanos Cancer Institute, Detroit, MI), and SUM149PT from Dr. S. Ethier (University of Michigan, Ann Arbor, MI). Identity of the cell lines was confirmed by short tandem repeats (STR) analysis. CAFs were derived from primary tumors and cultured in SUM medium as described before^56^. All cell lines were tested for mycoplasma infection routinely. Breast cancer cell lines: MDA-MB-231, SUM159PT, MDA-MB-453 were grown in McCoy medium (ThermoScientific) with 10% FBS (Life Technologies); T-47D, HCC1937 were grown in RPMI-1640 medium (ThermoScientific) with 10% FBS and 10 μg/ml human recombinant insulin (ThermoScientific); MCF10DCIS were grown in DMEM/F12 supplemented with 5% horse serum (ThermoScientific), 10 μg/ml human recombinant insulin, 20 ng/ml EGF (PeproTech), 100 ng/ml cholera toxin (ThermoFisher), 5 μg/ml hydrocortisone (Sigma); MCF7, HS578T were grown in DMEM/F12 with 10% FBS and 10 μg/ml human recombinant insulin, SUM149PT were grown in SUM medium (1:1 mix of DMEM/F12 and Human Mammary Epithelial Cell Growth Medium (Sigma), 5% FBS, 5 μg/ml). Fluorescently labelled derivates of carcinoma cell lines and fibroblasts were obtained by lentiviral expression of pLV[exp]-CAG-NLS-GFP-P2A-puro, pLV[exp]-CAG-NLS-mCherry-P2A-puro, pLV[Exp]-CAG>Bsd(ns):P2A:EGFP (custom vectors from VectorBuilder), or mCherry/Luciferase (obtained from Dr. C. Mitsiades, DFCI).

For fusion assays, 50/50 mixes of cells expressing differentially expressed fluorescent and antibiotic resistance markers were seeded into 6 cm or 10 cm tissue culture dishes (Sarstedt) in 50/50 mixture of DMEM-F12, 10% FBS/MEGM with supplements. Following co-culture for three days, the cells were harvested and subjected to flow cytometry analysis or replated for dual antibiotic selection with 10 μg/ml blasticidin and 2.5 μg/ml puromycin. Parental cells expressing single antibiotic resistance marker were used as negative controls. During the 7-14 days, required to eliminate all of the cells in the single resistance control plates, cells from mixed populations were replated once to relieve contact inhibition. After surviving cells from the mixed cultures approached confluency under double antibiotic selection, they were considered as passage 0. For each subsequent passage, 5×10^5^ cells cells were replated into 10 cm dishes and grown until ~90% confluency (5-7×10^6^ per dish). Antibiotic selection was maintained for these additional passages. For derivation of subclones, cells from mixed hybrid cultures were seeded into 96 well plate at seeding density <1 cell per well. Next day after seeding, the wells were examined and those containing >1 cells were excluded. Upon reaching >50 cells, the colonies were trypsinized and replated into 6 well plates, then into 10 cm dishes. Upon reaching confluency, the cells were divided for the DNA content and SNP inheritance analyses.

### Flow cytometry analyses

For the detection of hybrids, cells from monocultures or co-cultures of differentially labelled cells were harvested with 0.25% Trypsin (ThermoFisher) and resuspended in PBS with 0.1 μg/mL DAPI (Sigma). For the negative controls, cells from monocultures were mixed immediately after harvesting and kept on ice. Flow cytometry analyses were performed using MACSQuant VYB cytometer (Milteniy Biotec), with data analyzed using FlowJo software. The average number of collected events was 80,000. For the image-based flow cytometry analyses cells were incubated for 20 min in PBS with 5 μg/ml Hoechst33342 (ThermoFisher), prior to harvest. The analyses were performed with Amnis ImageStream X Mark II imaging flow cytometer (Amnis, Luminex), using IDEAS software (Amnis, Luminex). The average number of collected events was 10,000.

For the DNA content analyses 10^6^ of cells were resuspended in 500 μl of PBS and then 4.5 ml ice cold 70% ethanol was added dropwise. Cells were kept at −20C for at least 2 hours, then washed in PBS twice and resuspended in 300 μl PBS with 0.1% Triton X-100 (ThermoFisher), 0.2 mg/ml RNAse (Qiagen) and 20 μg/ml of propidium iodide (ThermoFisher). Flow cytometry analysis was performed using MACSQuant VYB instrument (Milteniy Biotec). For each of the tested pairs of co-cultures and controls, FACS analyses were performed over two or more distinct experiments.

### Microscopy studies

Live cell images were acquired with Axioscope microscope with A-Plan 10x/0.25 Ph1 objective and AxioCam ICm1 camera (Zeiss), using ZEN software (Zeiss). Time-lapse videos were generated with IncuCyte live cell imaging system (Sartorius) using ZOOM 10X objective for breast cancer cell mixes and ZOOM 4x objective for breast cancer cell-fibroblast mixes. Images were acquired in red and green fluorescent channels as well as visible light channel every 3 hours for 4-5 days. For immunofluorescent detection of GFP and mCherry in xenograft tumors, formalin fixed, paraffin embedded tumors were cut at 5 microns sections. Deparaffinized tissue slices were blocked in PBS with 10% goat serum for 30 min at room temperature, then incubated at room temperature for 1 hour with primary antibodies and 1 hour with secondary antibodies and 0.1 μg/mL DAPI (Sigma) with 3×10 min washes after each incubation. Vector TrueVIEW (Vector Labs) autofluorescence quenching reagent was used prior to mounting the slides. Anti dsRed (1:100, Clonetech #632496) was used for the detection of mCherry, anti-GFP 4B10 (1:100, CST #632496) was used for detection of GFP. Alexa Fluor Plus 647 goat anti-rabbit (1:1000, Invitrogen # A32733), and Alexa Fluor Plus 488 goat anti-mouse (1:1000, Invitrogen #A32723) were used as secondary antibodies. Confocal immunofluorescent images were acquired with Leica TCS SP5 system with 63x objective (Leica).

For histology analyses of metastatic outgrowth, formalin-fixed paraffin-embedded slides were sectioned at 5 microns, stained with hematoxylin and eosin and scanned with Aperio ScanScope XT Slide Scanner (Leica). Images were segmented into lung and metastatic regions. Segmentation and annotation of numbers and area of metastatic lesions were done using QuPath software (https://qupath.github.io/) in consultation with a pathologist.

### Mouse xenograft studies

To detect fusion *in vivo*, parental GFP and mCherry expressing cells were harvested and mixed at 50/50 ratio in DMEME-F12 culture media mixed with 50% Matrigel (BD Biosciences). The mixtures were injected into fat pads of 8 weeks old female NSG mice on both flanks, with 1×10^6^ cells in 100 μl volume per injection. Tumor growth was monitored weekly by palpation. When tumor diameter reached 1 cm or animals become moribund with symptoms of reduced mobility, hunching and labored breathing, mice were euthanized. Xenograft tumors were fixed in formaldehide and embedded in paraffin.

For the lung metastasis assays, luciferase-labelled parental cell lines (MCF10DCIS and SUM159PT) or their hybrids at passage 8 post antibiotic selection were harvested and resuspended in DMEM/F12 medium with 10% FBS. Approximately 2×10^5^ cells were injected per animal through the tail vein into 8-week-old NSG mice. Tumor growth was monitored by bioluminescence capture with the IVIS-200 imaging system (PerkinElmer) under isoflurane anesthesia. Imaging was performed immediately after cell injection and then weekly. One month after injection, mice were euthanized, and lungs were extracted. One lobe was fixed in 10% formalin for subsequent paraffin embedding and histology analysis while the remaining tissue was cut into small fragments and incubated in collagenase/hyaluronidase/BSA solution (5 mg/ml each) at 37°C under constant mixing, until disappearance of visible chunks of tissue. Following 2x wash with ice-cold PBS, cell suspension was filtered using 500 μm cell strainers (Puriselect), pelleted and plated into 10 cm dishes for a brief (3 days) *ex vivo* culture prior to FACS analyses. All animal experiments were performed in accordance with the guidelines of the IACUC of the H. Lee Moffitt Cancer Center.

### Growth and invasion/migration assays

To determine growth rates, 5×10^4^ cells were seeded in triplicates into 6 cm culture dishes. Upon reaching ~90% confluency, cells were harvested by trypsinization, counted and re-seeded at the same starting density. Growth rates were calculated as average ln(cell number fold change)/(number of days in culture) over three passages. Invasion/migration assay was performed in 12 well ThinCert plates with 8 μm pore inserts (Greiner Bio-One, #665638). Parental cells and hybrids were plated in appropriate FBS-free medium containing 0.1% BSA (Sigma) on transwell insert membrane covered with 30% Matrigel (BD Biosciences) in PBS. The lower wells contained medium with 10% FBS. Approximately 10^4^ cells were plated and cultured for 72 hours. Matrigel was removed from the transwell insert and cells migrated through the membrane fixed in 100% methanol for 10 min on ice and stained with 0.5% crystal violet solution in 25% methanol. Membranes were cut off and mounted on glass slides with xylene-based Cytoseal mounting media (ThermoFisher). Slides were scanned using Evos Autoimaging system (ThermoFisher), and images were analyzed with ImageJ software (https://fiji.sc/). Sixteen tiles per sample were analyzed.

### Colony formation assays

For *in vitro* clonogenic assays, 50/50 mixes of GFP- and mCherry-expressing parental cell lines were grown separately (controls) or in co-cultures for 3 days. For *ex vivo* clonogenic assays, tumors initiated from implantation of single-labelled cells or 50/50 mixes were harvested 4 weeks post implantation, and digested with the mixture of 2 mg/ml collagenase I (Worthington Biochem) and 2 mg/ml hyalurinidase (Sigma H3506) at 37°C for 3 hours. Approximately 1×10^6^ cells were seeded in 10 cm culture dishes in corresponding growth media with 10 μg/ml blasticidin and 2.5 μg/ml puromycin; for negative controls, 5×10^5^ of each of the parental cells were seeded. For clonogenicity rate controls, parental cells were seeded at 100 cells per 6 cm dish in corresponding antibiotic. Following 2 weeks of incubation, with selection media replaced twice per week and acquisition of fluorescent images of representative colonies, the medium was removed, plates were washed twice with PBS, and then colonies were fixed in a solution of 12.5% acetic acid and 30% methanol for 15 min and stained with 0.1% crystal violet solution in water for 4 hours. Colonies with approximately 50 or more cells were manually counted. Clonogenic frequency was calculated using numbers of colonies as a percentage of seeded cells. To account for reduced clonogenic survival of cells, freshly isolated from xenograft tumors, clonogenic data for the *ex vivo* samples was normalized to clonogenicity of *ex vivo* cells isolated tumors seeded with parental cells, plated in the absence of antibiotic selection.

### SNP and copy number analyses

DNA from ~3×10^6^ cells was extracted with DNeasy Blood&Tissue kit (Qiagen). For SUM159PT/MDA-MB-231 hybrids and MCFDCIS/SUM159PT from lung metastasis CytoSNP-12 v2.1 BeadChip array from Illumina was used and data analyzed with GenomeStudio 2.0 software (Illumina). For MCF10DCIS/SUM159PT and HS578T/MDA-MB-231 hybrids, CytoScan array from Affimetrix was used. SNP data analyzed and LogR Ratio plots were generated with ChAS software (Affimetrix). Data were analyzed and visualized using R (version 3.3.2). First, genotypes of parental cell lines were compared, and homozygous SNPs that are distinct between the two parental cell lines were selected. Then, for each of the differential SNP an identity score was assigned for a hybrid sample: 0 if only SNPs from the parent 1 were detected, 1 if only SNPs from parent 2 were detected, and 0.5 if both could be detected. The data were plotted as a heat map where rows represent hybrid samples and columns are aligned by position number of each SNP, with colors conveying the identity score. For copy number and genotype analyses and visualization of data from CytoScan and CytoSNP-12 SNP arrays, we used Chromosome Analysis Suite 4.1 (ChAS 4.1). We used Genome Studio to analyze the copy numbers and genotypes of the samples, respectively. Segmented copy numbers of the samples were visualized in the Integrative Genome Browser (IGV v2.6.2). Morpheus (https://software.broadinstitute.org/morpheus) online tool was used to visualize the differential allelic inheritance data.

### Single-cell RNA sequencing and analysis

Chromium Single Cell 3’ Library, Gel Bead & Multiplex Kit and Chip Kit (10X Genomics) was used to encapsulate and barcode for cDNA preparation of parental (SUM159PT and MDA-MB-231) and hybrid cells. Targeted cell population sizes were 2500 cells for each of the parental or hybrid samples. Libraries were constructed according to manufacturer’s protocol, sequenced on an Illumina NovaSeq, and mapped to the human genome (GRCh38) using CellRanger (10X Genomics) with an extended reference to include GFP and mCherry proteins that the parental lines were engineered to expressed.

Raw gene expression matrices generated per sample using CellRanger (version 3.0.1) were combined in R (version 3.5.2) and converted to a Seurat object using the Seurat R package (https://satijalab.org/seurat/)^57^. From this, all cells were removed that had over two-standard deviations of the mean UMIs derived from the mitochondrial genome (between 5-12% depending on sample) and only kept cells with gene expression within two-standard deviations of the mean gene expression^58^. From the remaining 10,059 gene expression matrices were log-normalized and scaled to remove variation due to total cellular read counts. To reduce dimensionality of this dataset, the first 200 principal components were calculated based on the top 2000 variable genes. Jack Straw analysis confirmed that the majority of the dataset variation were captured in these first principal components. All the cells were then clustered using the Louvain algorithm implemented by Seurat by creating a graph representation of the dataset with 10,059 nodes and 413,785 edges and optimizing based on modularity. With a resolution of 0.6, the algorithm identified 12 communities with a maximum modularity of 0.8931, which gave us confidence in this clustering ^59^. The data was then visualized by running the uniform manifold approximation and projection (UMAP) algorithm^28^ and either coloring individual cells based on their cluster identity or cell type identity. Gini coefficients were calculated with the package DescTools in R^60^. Zero values were excluded, and reads were used from 10x genomics outputs.

### Statistical analyses

Statistical analyses of *in vitro* and *in vivo* experimental data were performed using GraphPad Prism and Matlab software, using statistical tests indicated in figure legends.

### Computational methods

Muller’s plots were obtained using the RTool Evofreq^61^. The agent-based model was implemented in the JAVA framework HAL^62^. To visualize cells carrying unique genotypes (results presented in Fig. 5G and supplementary videos S6-9), the scalar value of the genotype binary vector value in base 2 was mapped to rainbow color scale. Detailed description of mathematical modeling is provided in the Mathematical Supplement.

## ACKNOWLEDGMENTS

We thank Dr. Aaron Goldman for sharing his unpublished observations of frequent occurrence of cell fusions in his experimental studies, which confirmed our observations and helped motivating this work. We thank Dr. Yuri Lazebnik, Dr. Aaron Goldman and Dr. Doris Tabassum for providing thoughtful feedback. We thank Rafael Bravo for the help he provided for using the framework HAL We thank the Flow Cytometry, Analytic Microscopy, Tissue Histology, Biostatistics and Bioinformatics Shared Resource and Molecular Genomic Core Facilities at the H. Lee Moffitt Cancer Center & Research Institute; an NCI designated Comprehensive Cancer Center (P30-CA076292). We thank Moffitt SPARK program for supporting internship for MAL. This work was supported by Susan G. Komen Breast Cancer Foundation CCR17481976 (A.M.), Moffitt Cancer Biology and Evolution program pilot award (AM), and Integrative Mathematical Oncology Workshop award (AM and DB).

---

**S1.**
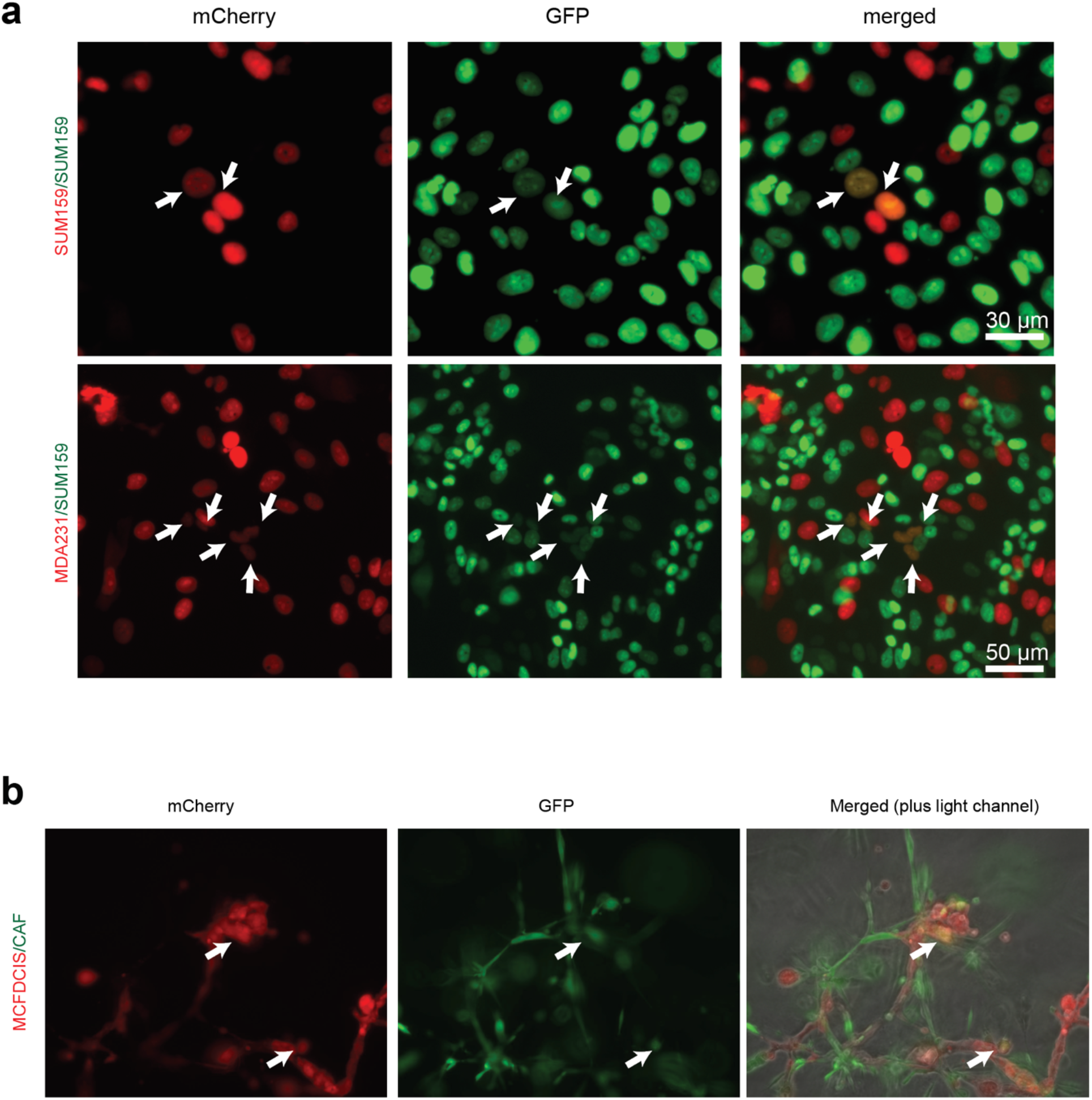
**a.** Detection of putative spontaneous cell fusions. Live-cell fluorescence microscopy images of 2D co-cultures between differentially labelled (nuclear GFP and mCherry) cell lines. Arrowheads indicate cells that express both labels. **b.** Live-cell fluorescence microscopy images of 3D Matrigel co-cultures between MCF10DCIS breast carcinoma cells and a primary breast CAF isolate labelled with cytoplasmic GFP and dsRED, respectively. Arrowheads indicate cells that express both labels.

**S2.**
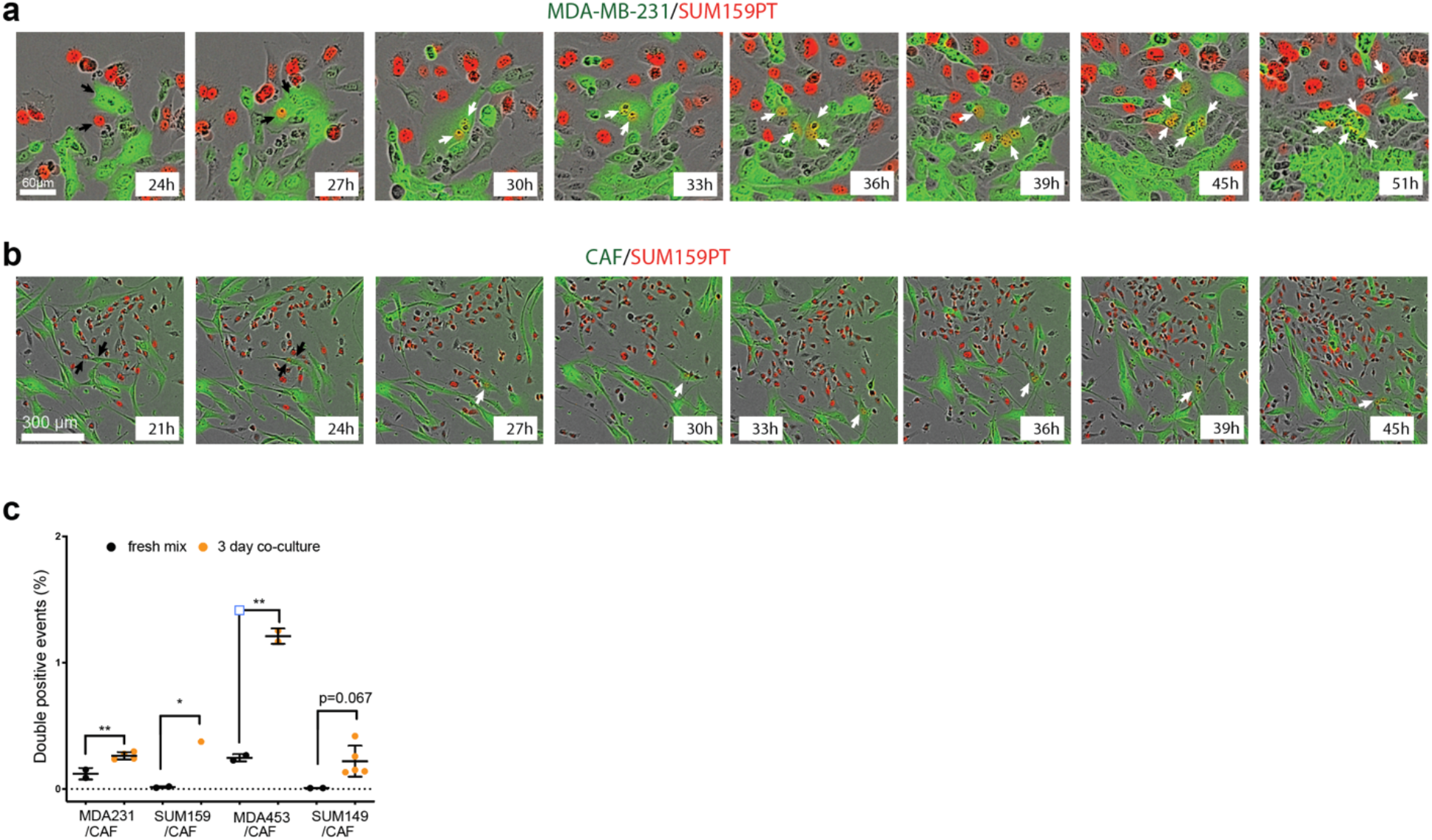
Time-lapse live fluorescence microscopy images of mCherry+ SUM159PT cells co-cultured with GFP+ MDA-MB-231 cells (**a**) and CAFs (**b**). The labels indicate time after plating. Black arrowheads show fusion parents, and white arrowheads show double-positive hybrid cells and their progeny. **c.** Quantification of flow cytometry detection of double-positive events in the cocultures between GFP+ CAFs and indicated breast cancer cell lines labelled with mCherry. *and ** denote p values below 0.05 and 0.01 for two-tailed unpaired t-test, respectively.

**S3.**
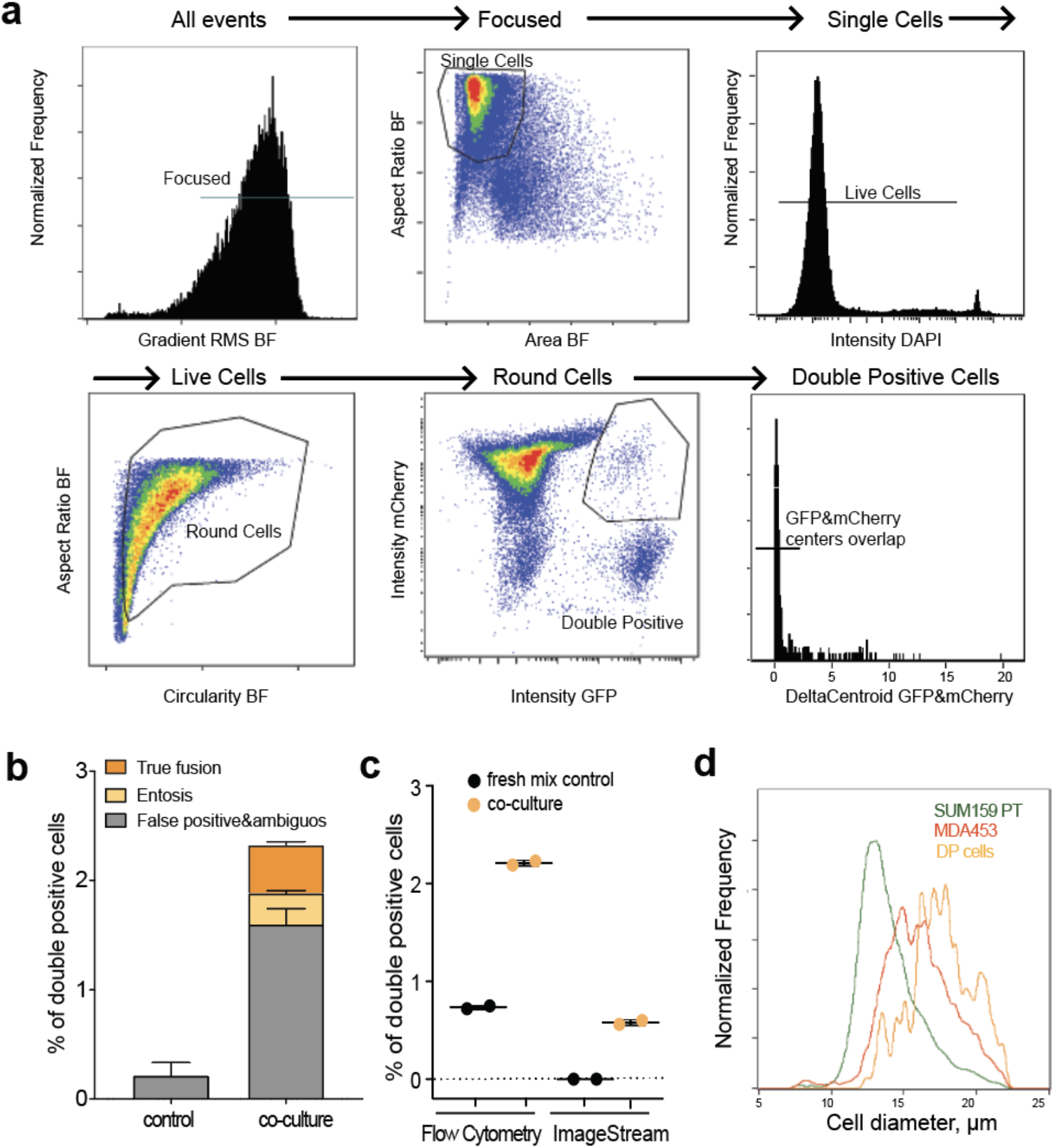
ImageStream detection of spontaneous cell fusions. **a.** Gating strategy for the detection of double-positive cells with ImageStream imaging-based flow cytometry platform. **b.** Quantitation of different classes of double-positive events in 3-day co-cultures between 50/50 mixes of GFP/mCherry labelled MCF7 cells, with examples of diffedrent classes of events provided in **Fig. 1e**. **c**. Comparison of frequency of double-positive events detected from the same samples of co-cultures of differentially labelled MDA-MB-231 (mCherry+) and SUM159PT (GFP+) cells and freshly mixed controls using FACS and ImageStream platforms (validated true positives percentages are plotted for ImageStream analyses). **d.** Distribution of cell diameters of the parental and double-positive cells from ImageStream data shown in (**c**) measured in bright field and plotted using IDEAS software (ImageStream).

**S4.**
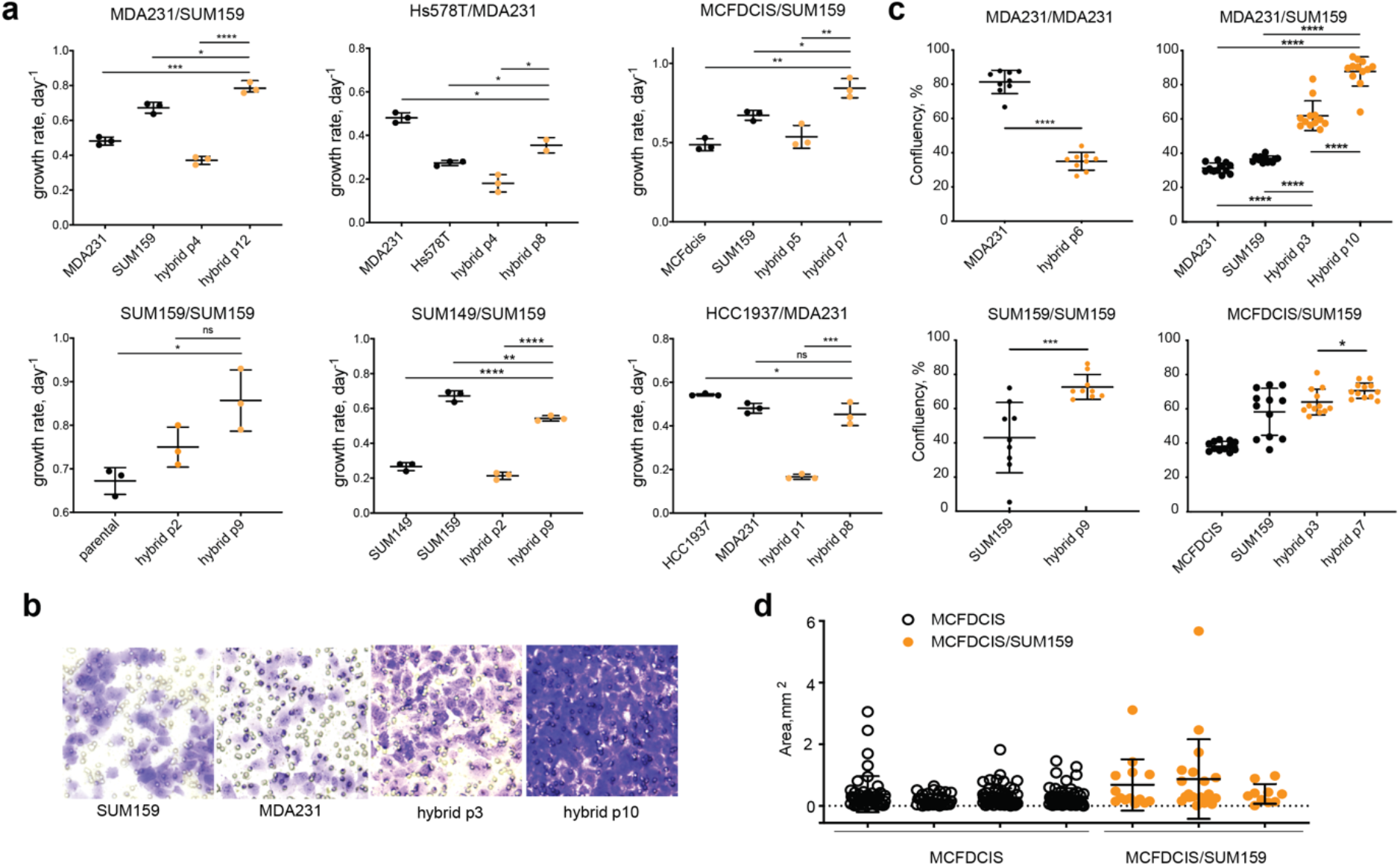
Phenotypic characterization of hybrid cells. **a.** Growth rates of the indicated cell lines and their hybrids, at the indicated passages post-antibiotic-selection. **b.** Representative images of stained membranes from Boyden chamber cell invasion/migration assay. **c**. Quantitation of Boyden chamber cell invasion/migration assay data. **d.** Quantification of area of lung metastases formed after tail vein injection of MCF10DCIS and MCF10DCIS/SUM159PT hybrids. Data from individual mice are plotted separately. *, **, ***, *** denote p values below 0.05, 0.01, 0.001 and 0.0001, respectively, for the two-tailed unpaired t-test.

**S5.**
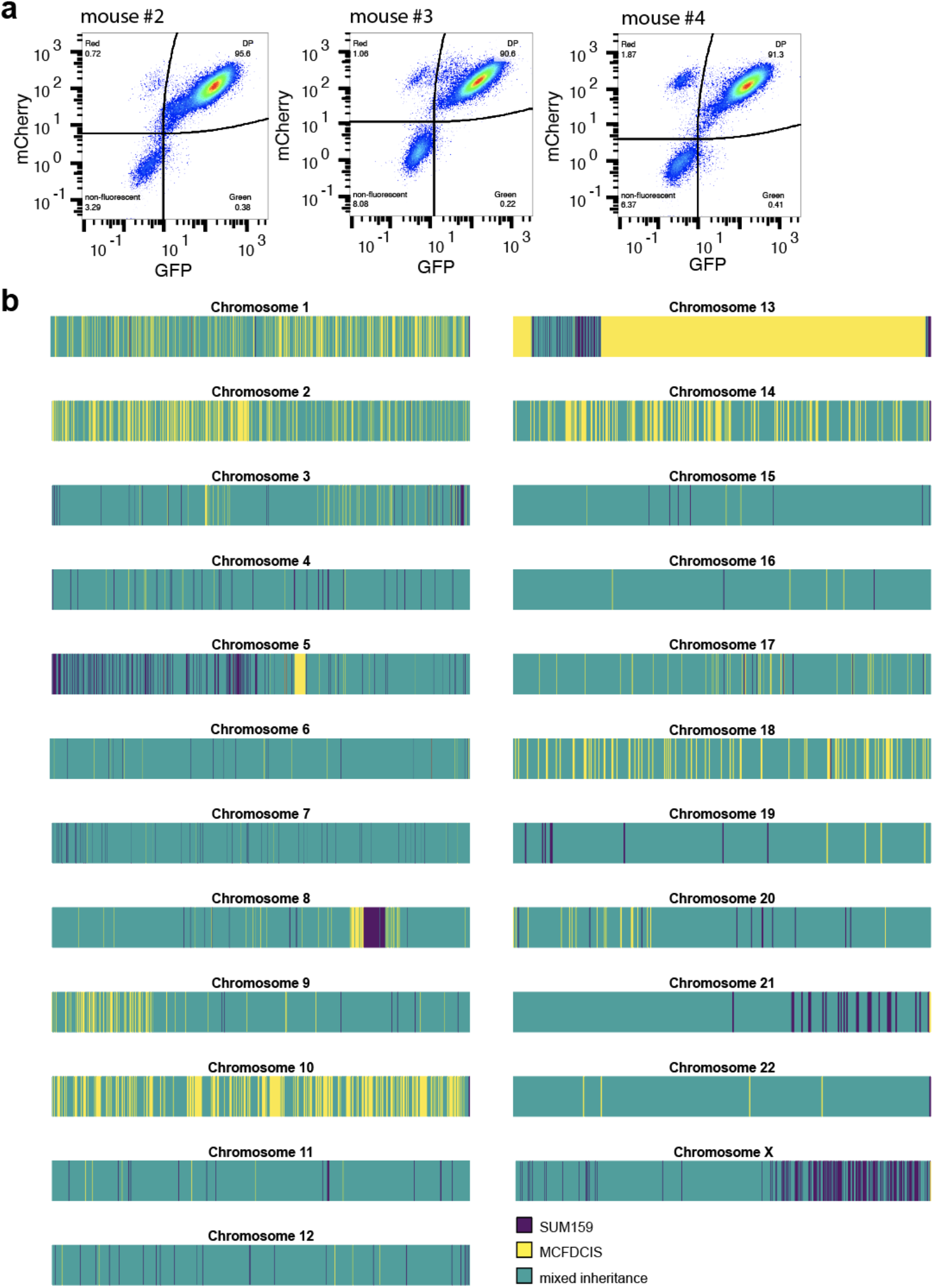
Inheritance of parent specific SNP alleles in lung tumors seeded with SUM159PT /MCFDCIS/ hybrids. **a.** FACS analysis of retention of GFP and mCherry fluorescence in hybrid cells, recovered from metastasis-bearing lungs of the indicated animals depicted in Fig. 2g. **b**. Mapping of parental cell line-specific SNP alleles (detected with Illumina CytoSPN-12 platform) across the genome of the hybrid cells isolated from mouse #3. Each chromosome is depicted in a separate panel. Columns indicate parent-specific alleles, as described in the color key.

**S6.**
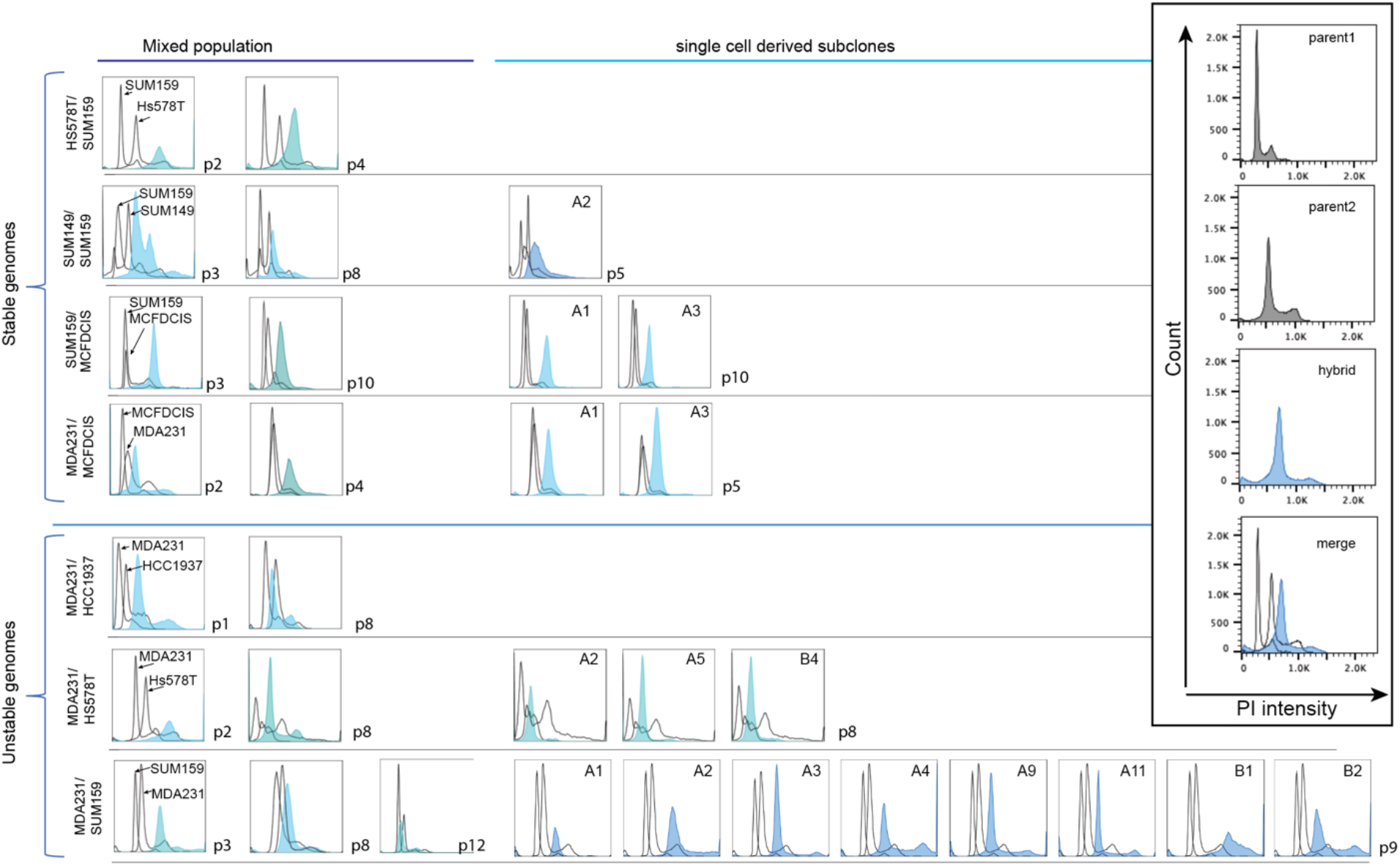
DNA content analysis of hybrid cells. FACS analysis of DNA content of the indicated parental cell lines and their hybrids. P1-12 indicate passage number of the mixed hybrid populations; for hybrid subclones the number indicates passage of the mixed culture, used for isolation of single cell subclones. Black contours indicate DNA content profiles for parental cell lines and their G1 and G2 peaks are used as reference points, filled histograms indicate DNA content profiles of the hybrids. “Unstable genomes” and “stable genomes” refer to hybrids that, respectively, did and did not show evidence of reduced DNA content between early and extended passages. Hybrids with stable genomes demonstrate G1 peak closer to G2 peak of parents and no observed difference in DNA content between passages and single cell clones. Inset indicates axes and shows components of the merged plots.

**S7.**
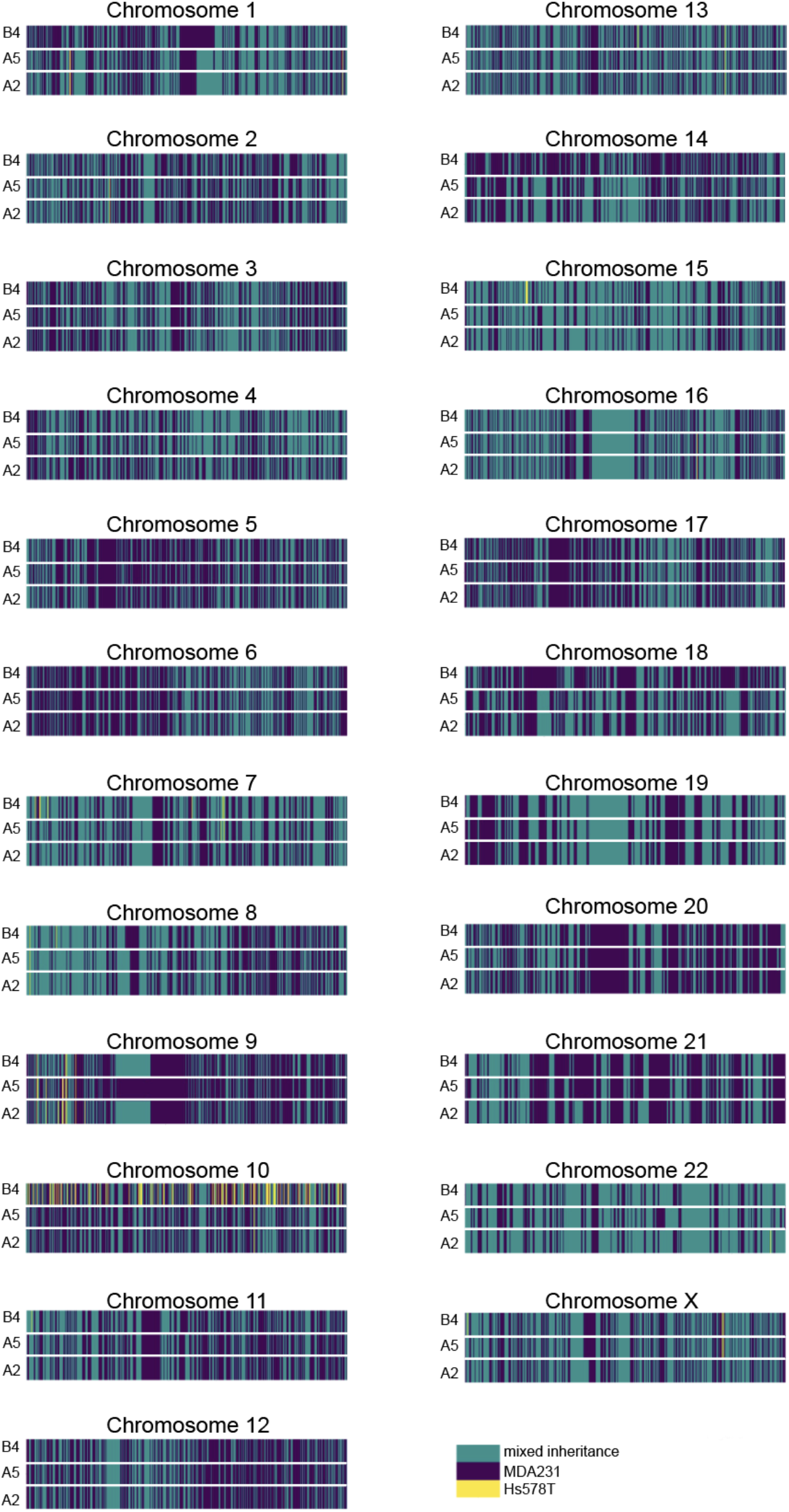
Allelic inheritance analyses of cell line-specific alleles for MDA-MB-231/HS578T/ hybrids. SNP inheritance data was obtained with Affymetrix CytoScan SNP array. Each chromosome is depicted in a separate panel. Rows depict distinct subclones shown in Fig. S6; columns indicate inheritance of parent-specific SNP alleles, as described in the color key.

**S8.**
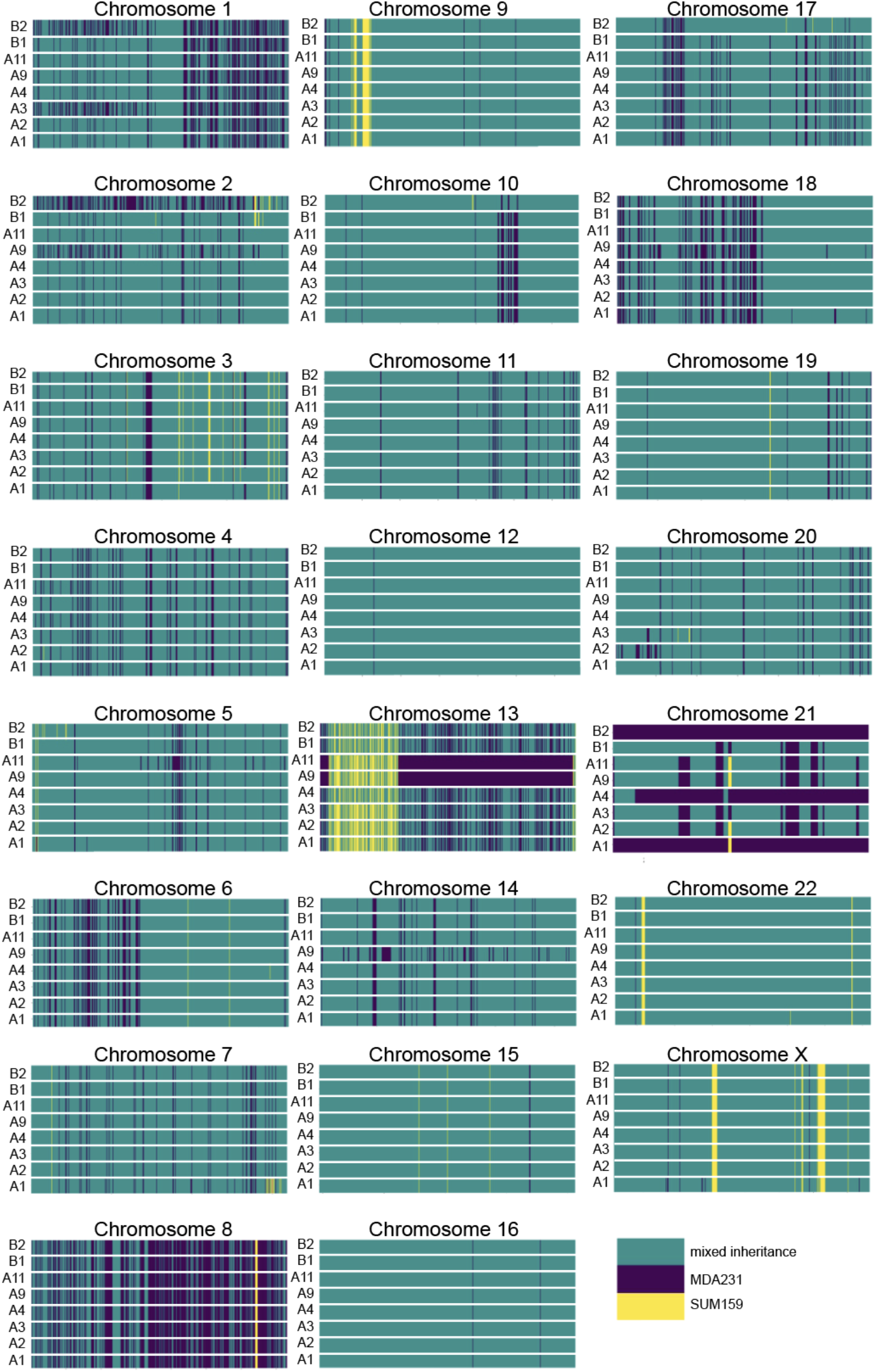
Allelic inheritance analyses of cell line-specific alleles for MDA-MB-231/SUM159PT hybrids. SNP inheritance data was obtained with Illumina CytoSPN-12 platform. Each chromosome is depicted in a separate panel. Rows depict distinct subclones shown in Fig. S6; columns indicate inheritance of parent-specific SNP alleles, as described in the color key.

**S9.**
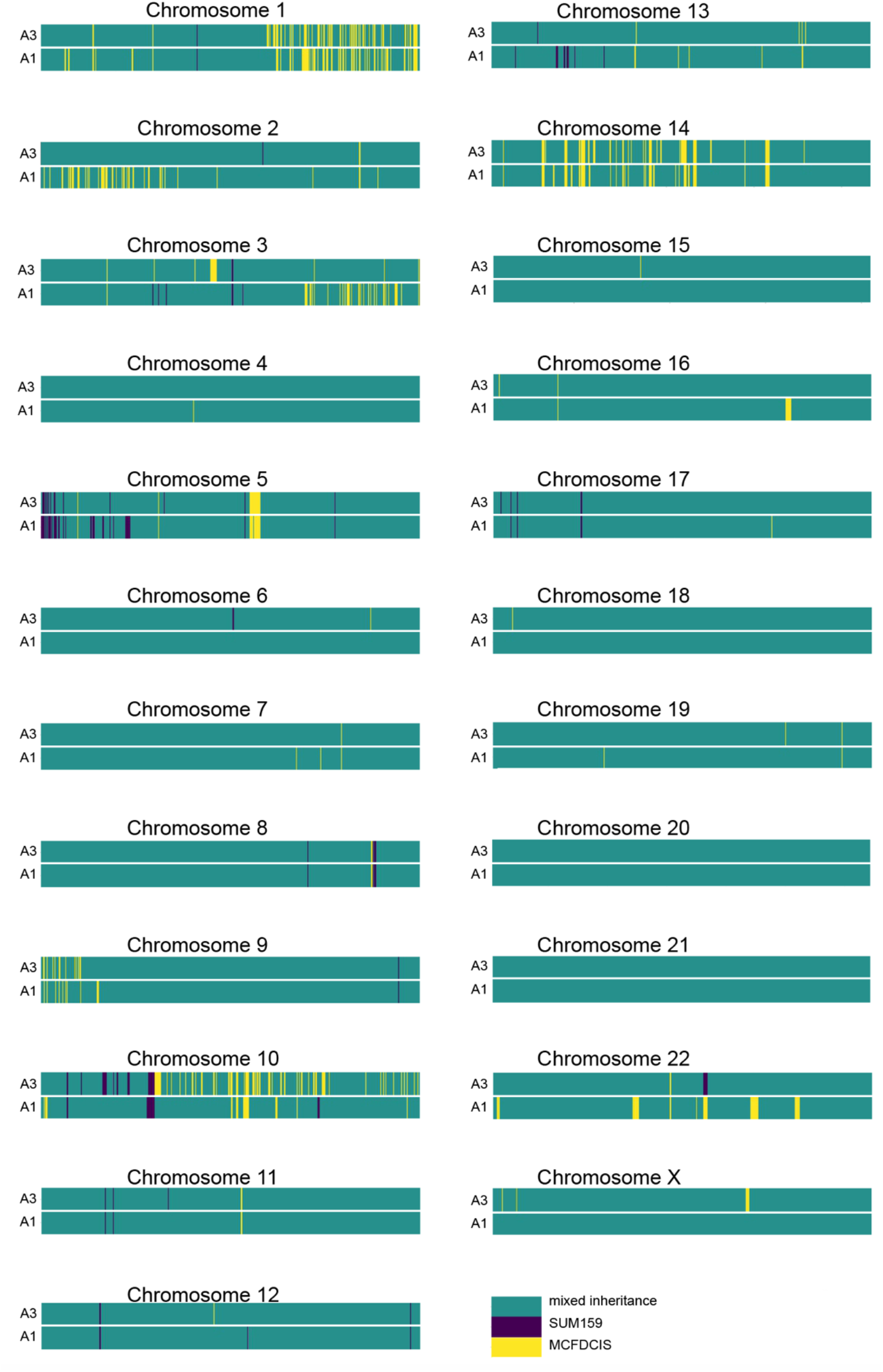
Allelic inheritance analyses of cell line-specific alleles for SUM159PT/MCF10DCIS hybrids. SNP inheritance data was obtained with Illumina CytoSPN-12 platform. Each chromosome is depicted in a separate panel. Rows depict distinct subclones shown in Fig. S5; columns indicate inheritance of parent-specific SNP alleles, as described in the color key.

**S10.**
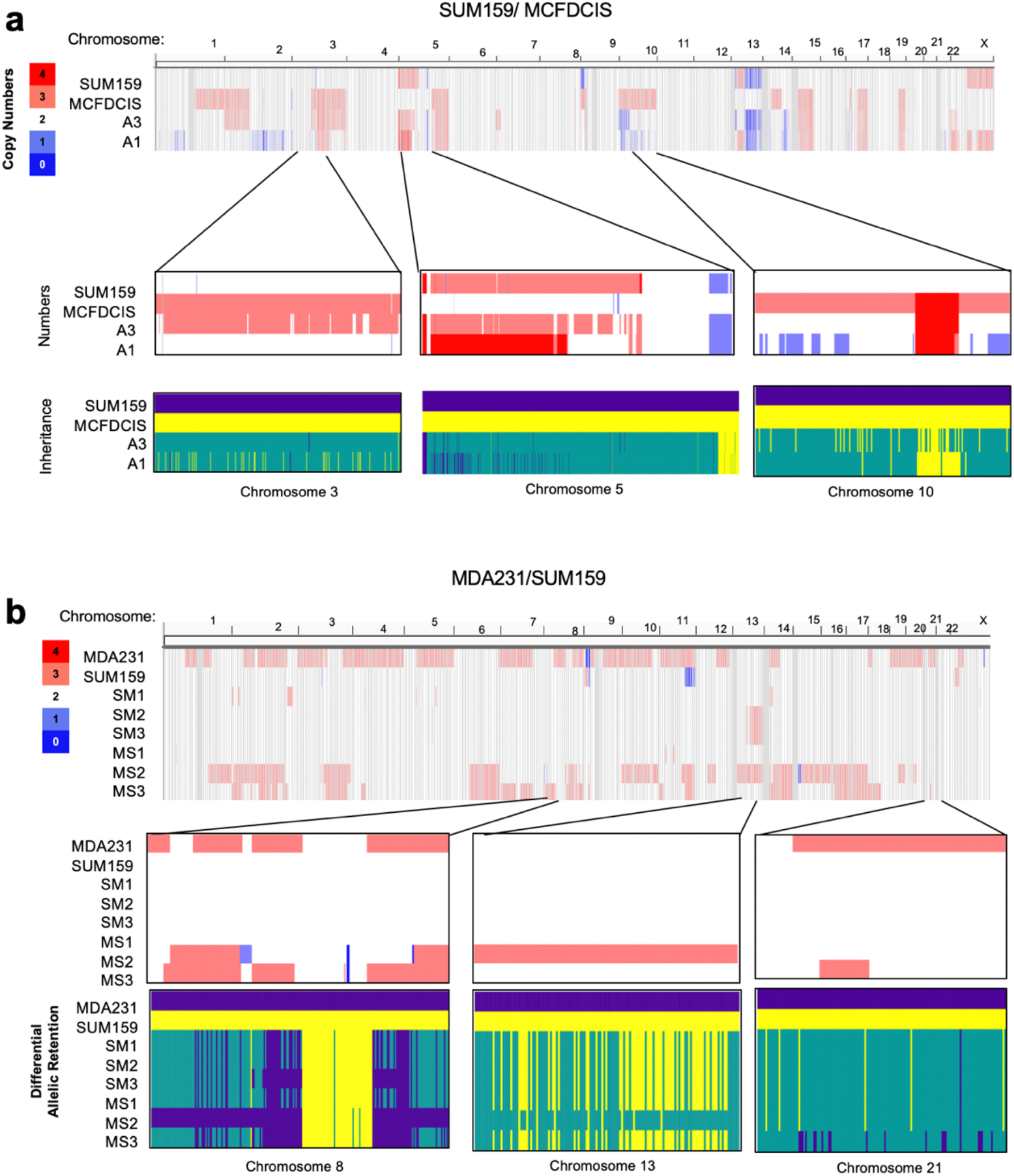
Copy number analyses of cell line-specific SNP alleles. SNP copy number and inheritance data were obtained with Illumina CytoSPN-12 platform. Copy number data status is shown for all of the cell line-specific SNP alleles across the genome for the indicated sublones. For the indicated selected zoom-in insets, correspondence between SNP copy numbers and detected inheritance is plotted. Turquoise color indicates mixed inheritance. **a**. Analyses of the SUM159/DCIS hybrids shown in Fig. 3g and Fig. S9. b. Analyses of subclones from MDA-MB-231/SUM159 hybrids, seeded from 1-t passage. SM1-3 and MS1-3 represent subclones isolated from two distinct hybrid populations, using opposite fluorophore/antibiotic labels of the parental cells.

**S11.**
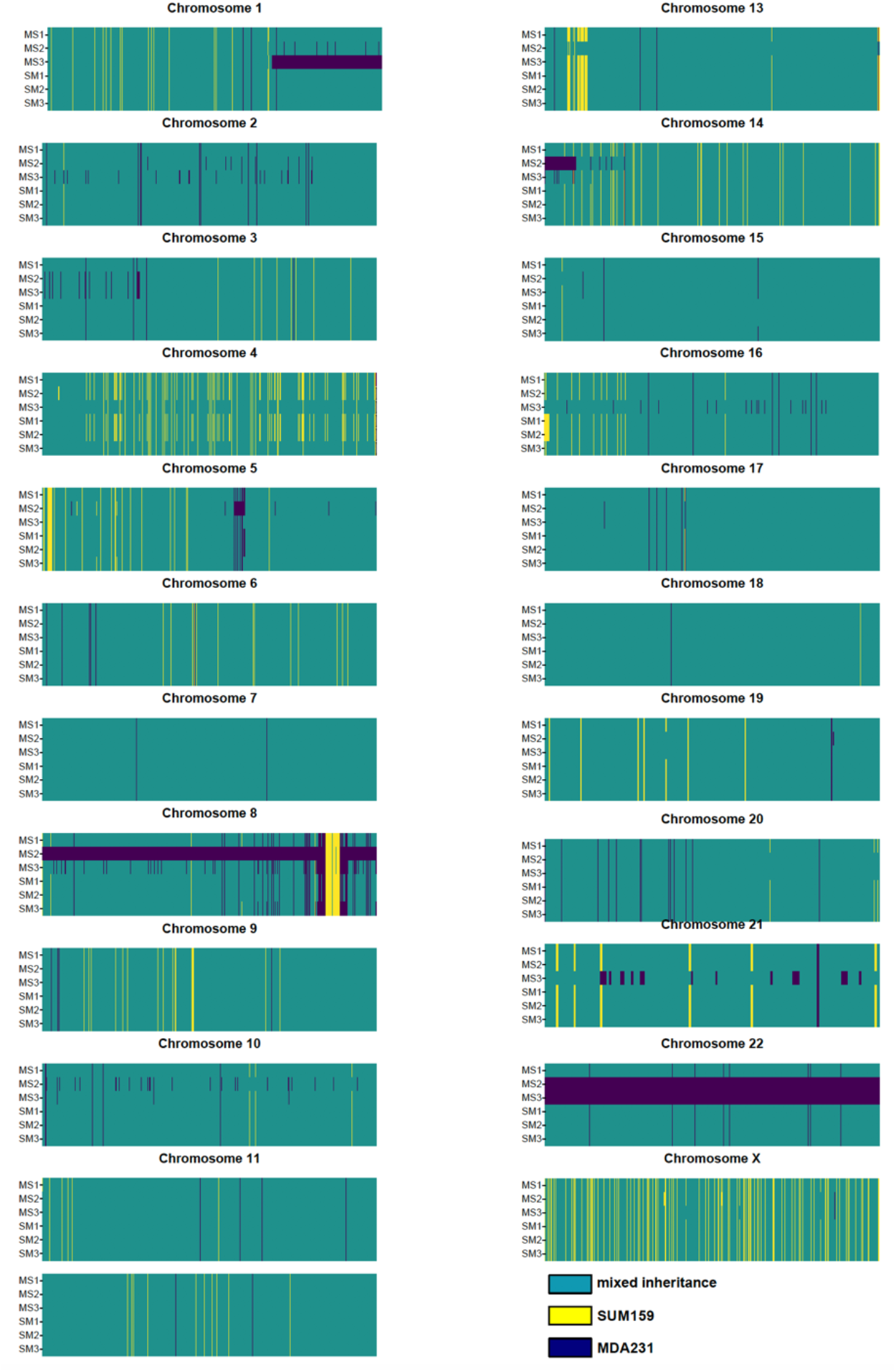
Allelic inheritance analyses of cell line-specific alleles for independently derived sets of MDA-MB-231/SUM159PT hybrids. SNP inheritance data was obtained with Illumina CytoSPN-12 platform. SM1-3 and MS1-3 represent subclones isolated from distinct hybrid populations. Lower degree of genetic divergence, compared to the hybrids shown in Fig. S 8 likely reflects earlier passage of subclone isolation seeding (passage 1 *vs*. passage 9).

**S12.**
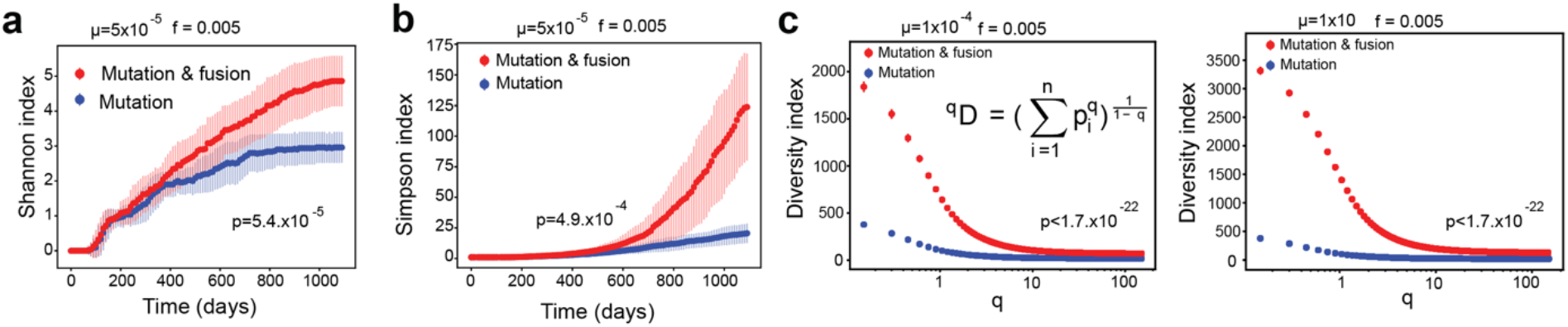
Impact of fusions on diversification. Comparisons are drawn between results of *in silico* simulations involving mutations only versus mutation and fusion. Clonal diversity is captured by Shannon (**a**), Simpson (**b**) and GDI (**c**) diversity indexes. Mutation and fusion rates are indicated in the figures. Indicated p values denote the results for Kolmogorov-Smirnov test for the final timepoint of simulations (a, b), or for the all of the <1 q values (c, d).

**S13.**
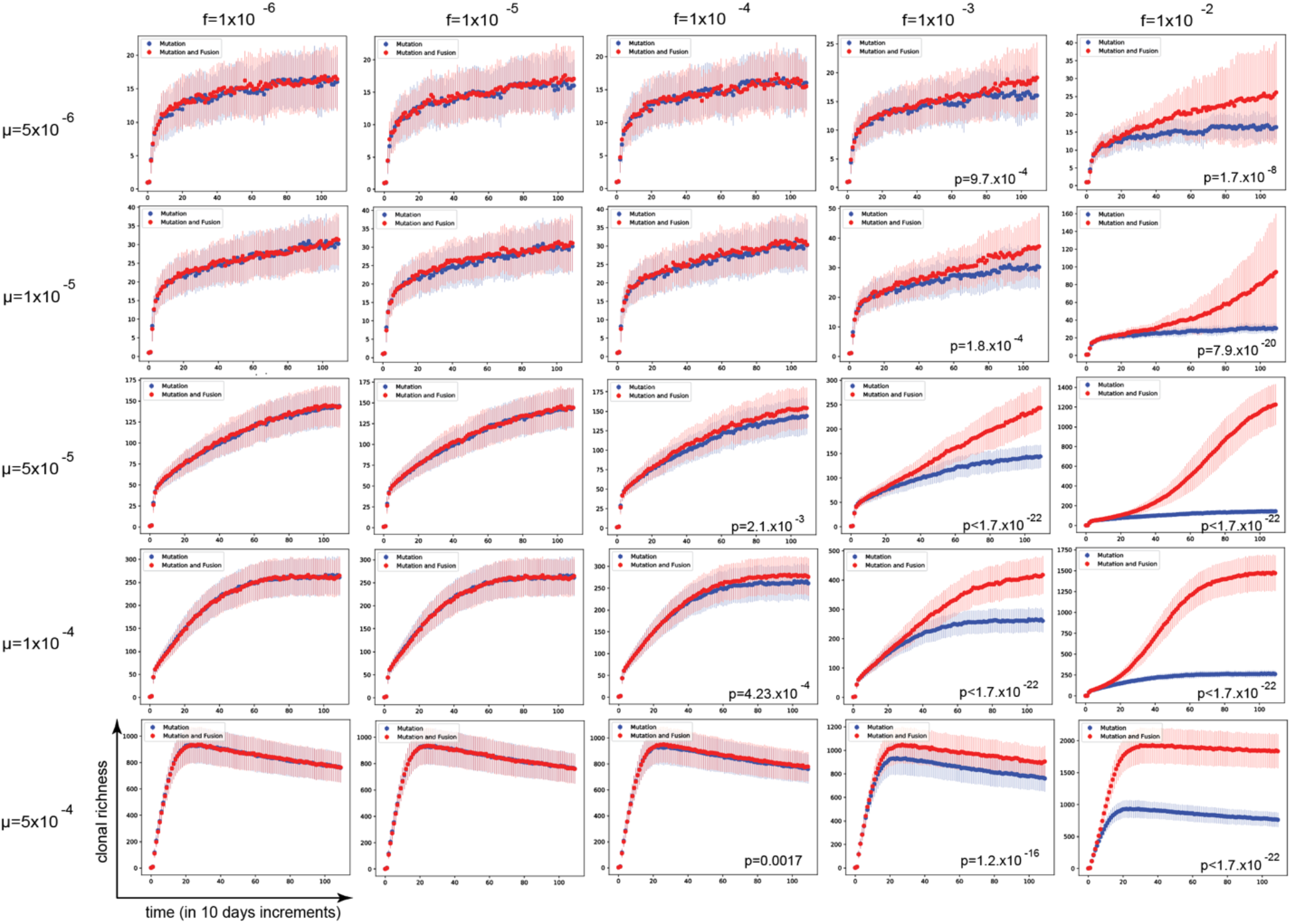
Parameter sweep analysis for the impact of mutation and fusion rates on clonal richness. Graphs depict results of *in silico* simulations with branching birth-death model showing clonal richness over time for the indicated mutation (μ) and fusion (f) rates; p values indicate the results of Kolmogorov-Smirnov test comparing clonal richness at the end of the simulation (only shown for differences that has reached the 0.05 significance threshold).

**S14.**
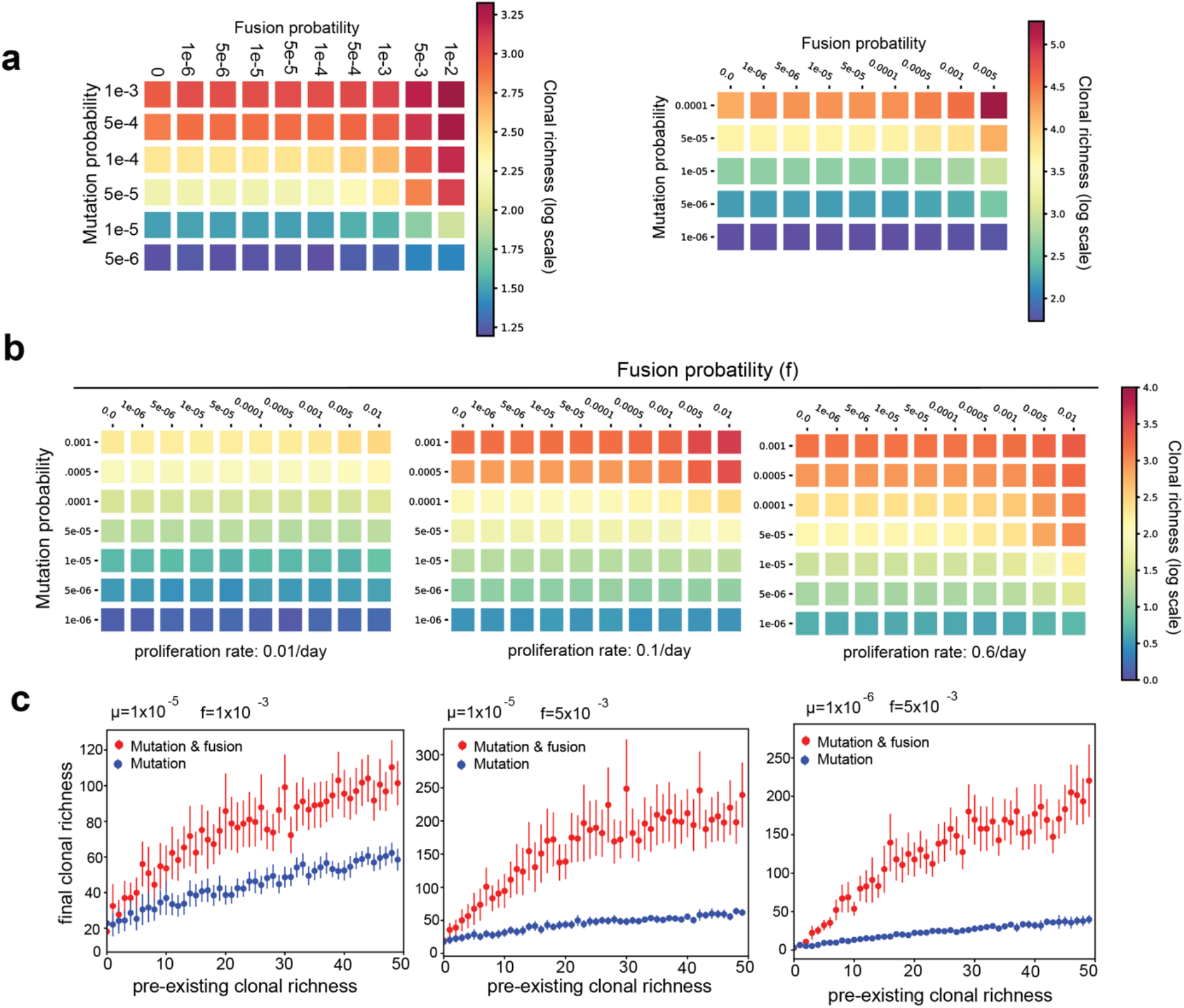
Impact of population size, cell turnover and starting genetic heterogeneity on diversification by fusion-mediated recombination. **a**. Exploration of the impact of maximal tumor population size (carrying capacity) on diversification over indicated mutation and fusion rates. **b**. Exploration of the impact of tumor proliferation/turnover rates on diversification over indicated mutation and fusion rates. **c**. Relationship between the initial clonal richness, and clonal richness at the indicated mutation and fusion rates. In all the panels, clonal richness at the end of the 1095 days of *in silico* growth is depicted.

**S15.**
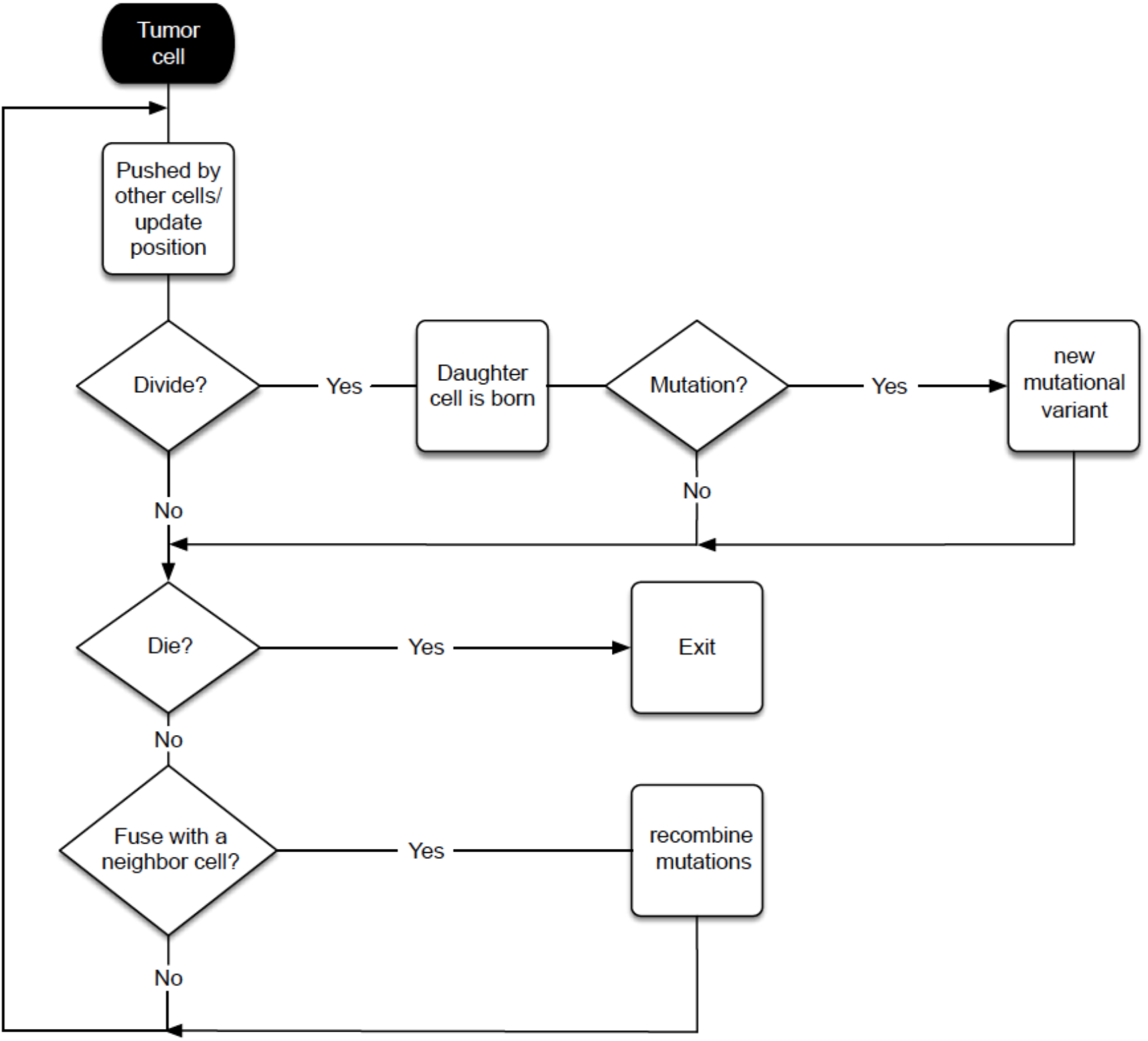
Flowchart for agent based model for spatial simulations.

## Supplementary Tables

**S1**. Genome-wide inheritance of parental specific SNP alleles in hybrid subclones.

**S2.** *In vitro* fusion rates estimated by ordinary differential equation data fitting to *in vitro* data. Rates were inferred with and without proliferation of hybrids. Standard deviation values of the parameter estimations are provided.

**S3.** *In vivo* fusion rates estimated by ordinary differential equation data fitting to *in vivo* data. Rates were inferred with either Logistic or Gompertz tumor growth models. Standard deviation values of the parameter estimations are provided.

## Supplementary Videos

**S1, S2.** Time lapse videos of co-cultures between MDA-MB-231cells labelled with cytoplasmic GFP and nuclear mCherry. Images were acquired every 3 hours over 5 days.

**S3, S4.** Time lapse videos of co-cultures between MDA-MB-231cells labelled with cytoplasmic GFP and SUM159PT cells labelled with nuclear mCherry. Images were acquired every 3 hours over 5 days.

**S5.** Time lapse videos of co-cultures between SUM159PT labelled with nuclear mCherry and CAFS labelled with nuclear GFP. co-cultured with CAFs with nuclear GFP. Images were acquired every 3 hours over 5 days.

**S6** Video of the entire simulation corresponding to Fig5G (2D mutation only).

**S7** Video of the entire simulation corresponding to Fig5G (2D mutation and fusion)

**S8** Video of the entire simulation corresponding to Fig5G (3D mutation only)

**S9** Video of the entire simulation corresponding to Fig5G (3D mutation and fusion)

**S10** Video of 3 days *in vitro* spatial simulation for the inferences of fusion rates described in the Mathematical Supplement.

## CITED REFERENCES

1 Greaves, M. & Maley, C. C. Clonal evolution in cancer. Nature 481, 306–313, doi:10.1038/nature10762 (2012).

2 Scott, J. & Marusyk, A. Somatic clonal evolution: A selection-centric perspective. Biochim Biophys Acta Rev Cancer 1867, 139–150, doi:10.1016/j.bbcan.2017.01.006 (2017).

3 Hanahan, D. & Weinberg, R. A. Hallmarks of cancer: the next generation. Cell 144, 646–674, doi:10.1016/j.cell.2011.02.013 (2011).

4 Becks, L. & Agrawal, A. F. The evolution of sex is favoured during adaptation to new environments. PLoS biology 10, e1001317, doi:10.1371/journal.pbio.1001317 (2012).

5 McDonald, M. J., Rice, D. P. & Desai, M. M. Sex speeds adaptation by altering the dynamics of molecular evolution. Nature 531, 233–236, doi:10.1038/nature17143 (2016).

6 Sinai, S., Olejarz, J., Neagu, I. A. & Nowak, M. A. Primordial sex facilitates the emergence of evolution. J R Soc Interface 15, doi:10.1098/rsif.2018.0003 (2018).

7 Duelli, D. & Lazebnik, Y. Cell fusion: a hidden enemy? Cancer cell 3, 445–448 (2003).

8 Lu, X. & Kang, Y. Cell fusion as a hidden force in tumor progression. Cancer research 69, 8536–8539, doi:10.1158/0008-5472.CAN-09-2159 (2009).

9 Platt, J. L., Zhou, X., Lefferts, A. R. & Cascalho, M. Cell Fusion in the War on Cancer: A Perspective on the Inception of Malignancy. Int J Mol Sci 17, doi:10.3390/ijms17071118 (2016).

10 Kuznetsova, A. Y. et al. Chromosomal instability, tolerance of mitotic errors and multidrug resistance are promoted by tetraploidization in human cells. Cell Cycle 14, 2810–2820, doi:10.1080/15384101.2015.1068482 (2015).

11 Fujiwara, T. et al. Cytokinesis failure generating tetraploids promotes tumorigenesis in p53-null cells. Nature 437, 1043–1047, doi:10.1038/nature04217 (2005).

12 Su, Y. et al. Somatic Cell Fusions Reveal Extensive Heterogeneity in Basal-like Breast Cancer. Cell Rep 11, 1549–1563, doi:10.1016/j.celrep.2015.05.011 (2015).

13 Zhou, X. et al. Cell Fusion Connects Oncogenesis with Tumor Evolution. Am J Pathol 185, 2049–2060, doi:10.1016/j.ajpath.2015.03.014 (2015).

14 Erenpreisa, J. & Cragg, M. S. MOS, aneuploidy and the ploidy cycle of cancer cells. Oncogene 29, 5447–5451, doi:10.1038/onc.2010.310 (2010).

15 Bennett, R. J. The parasexual lifestyle of Candida albicans. Curr Opin Microbiol 28, 10–17, doi:10.1016/j.mib.2015.06.017 (2015).

16 Zuba-Surma, E. K., Kucia, M., Abdel-Latif, A., Lillard, J. W., Jr. & Ratajczak, M. Z. The ImageStream System: a key step to a new era in imaging. Folia Histochem Cytobiol 45, 279–290 (2007).

17 Fais, S. & Overholtzer, M. Cell-in-cell phenomena in cancer. Nature reviews. Cancer 18, 758–766, doi:10.1038/s41568-018-0073-9 (2018).

18 Rappa, G., Mercapide, J. & Lorico, A. Spontaneous formation of tumorigenic hybrids between breast cancer and multipotent stromal cells is a source of tumor heterogeneity. Am J Pathol 180, 2504–2515, doi:10.1016/j.ajpath.2012.02.020 (2012).

19 Gast, C. E. et al. Cell fusion potentiates tumor heterogeneity and reveals circulating hybrid cells that correlate with stage and survival. Sci Adv 4, eaat7828, doi:10.1126/sciadv.aat7828 (2018).

20 Lu, X. & Kang, Y. Efficient acquisition of dual metastasis organotropism to bone and lung through stable spontaneous fusion between MDA-MB-231 variants. Proc Natl Acad Sci U S A 106, 9385–9390, doi:10.1073/pnas.0900108106 (2009).

21 Duelli, D. M. et al. A virus causes cancer by inducing massive chromosomal instability through cell fusion. Curr Biol 17, 431–437, doi:10.1016/j.cub.2007.01.049 (2007).

22 Storchova, Z. & Pellman, D. From polyploidy to aneuploidy, genome instability and cancer. Nat Rev Mol Cell Biol 5, 45–54, doi:10.1038/nrm1276 (2004).

23 Forche, A. et al. The parasexual cycle in Candida albicans provides an alternative pathway to meiosis for the formation of recombinant strains. PLoS biology 6, e110, doi:10.1371/journal.pbio.0060110 (2008).

24 Duncan, A. W. et al. The ploidy conveyor of mature hepatocytes as a source of genetic variation. Nature 467, 707–710, doi:10.1038/nature09414 (2010).

25 Skinner, A. M., Grompe, M. & Kurre, P. Intra-hematopoietic cell fusion as a source of somatic variation in the hematopoietic system. J Cell Sci 125, 2837–2843, doi:10.1242/jcs.100123 (2012).

26 Marusyk, A., Janiszewska, M. & Polyak, K. Intratumor Heterogeneity: The Rosetta Stone of Therapy Resistance. Cancer cell 37, 471–484, doi:10.1016/j.ccell.2020.03.007 (2020).

27 Koulakov, A. A. & Lazebnik, Y. The problem of colliding networks and its relation to cell fusion and cancer. Biophys J 103, 2011–2020, doi:10.1016/j.bpj.2012.08.062 (2012).

28 Becht, E. et al. Dimensionality reduction for visualizing single-cell data using UMAP. Nat Biotechnol, doi:10.1038/nbt.4314 (2018).

29 Ferrall-Fairbanks, M. C., Ball, M., Padron, E. & Altrock, P. M. Leveraging Single-Cell RNA Sequencing Experiments to Model Intratumor Heterogeneity. JCO Clin Cancer Inform 3, 1–10, doi:10.1200/CCI.18.00074 (2019).

30 Hill, M. O. Diversity and Evenness: A Unifying Notation and Its Consequences. Ecology 54, 427–432, doi:10.2307/1934352 (1973).

31 Rosenzweig, M. L., L, R. M. & Press, C. U. Species Diversity in Space and Time. (Cambridge University Press, 1995).

32 Hinohara, K. et al. KDM5 Histone Demethylase Activity Links Cellular Transcriptomic Heterogeneity to Therapeutic Resistance. Cancer cell 35, 330–332, doi:10.1016/j.ccell.2019.01.012 (2019).

33 Loeb, L. A. Human Cancers Express a Mutator Phenotype: Hypothesis, Origin, and Consequences. Cancer research 76, 2057–2059, doi:10.1158/0008-5472.CAN-16-0794 (2016).

34 Waclaw, B. et al. A spatial model predicts that dispersal and cell turnover limit intratumour heterogeneity. Nature 525, 261–264, doi:10.1038/nature14971 (2015).

35 Kimmel, G. J., Gerlee, P. & Altrock, P. M. Time scales and wave formation in non-linear spatial public goods games. PLoS Comput Biol 15, e1007361, doi:10.1371/journal.pcbi.1007361 (2019).

36 Gallaher, J. A., Enriquez-Navas, P. M., Luddy, K. A., Gatenby, R. A. & Anderson, A. R. A. Spatial Heterogeneity and Evolutionary Dynamics Modulate Time to Recurrence in Continuous and Adaptive Cancer Therapies. Cancer research 78, 2127–2139, doi:10.1158/0008-5472.CAN-17-2649 (2018).

37 Noble, R., Burri, D., Kather, J. N. & Beerenwinkel, N. Spatial structure governs the mode of tumour evolution. bioRxiv, 586735, doi:10.1101/586735 (2019).

38 Jacobsen, B. M. et al. Spontaneous fusion with, and transformation of mouse stroma by, malignant human breast cancer epithelium. Cancer research 66, 8274–8279, doi:10.1158/0008-5472.CAN-06-1456 (2006).

39 Mortensen, K., Lichtenberg, J., Thomsen, P. D. & Larsson, L. I. Spontaneous fusion between cancer cells and endothelial cells. Cell Mol Life Sci 61, 2125–2131, doi:10.1007/s00018-004-4200-2 (2004).

40 Melzer, C., von der Ohe, J. & Hass, R. In Vivo Cell Fusion between Mesenchymal Stroma/Stem-Like Cells and Breast Cancer Cells. Cancers 11, doi:10.3390/cancers11020185 (2019).

41 Searles, S. C., Santosa, E. K. & Bui, J. D. Cell-cell fusion as a mechanism of DNA exchange in cancer. Oncotarget 9, 6156–6173, doi:10.18632/oncotarget.23715 (2018).

42 Zack, T. I. et al. Pan-cancer patterns of somatic copy number alteration. Nat Genet 45, 1134–1140, doi:10.1038/ng.2760 (2013).

43 Dewhurst, S. M. et al. Tolerance of whole-genome doubling propagates chromosomal instability and accelerates cancer genome evolution. Cancer Discov 4, 175–185, doi:10.1158/2159-8290.CD-13-0285 (2014).

44 Stephens, P. J. et al. Massive genomic rearrangement acquired in a single catastrophic event during cancer development. Cell 144, 27–40, doi:10.1016/j.cell.2010.11.055 (2011).

45 Birdsell, J. A. & Wills, C. in Evolutionary Biology (eds Ross J. Macintyre & Michael T. Clegg) 27-138 (Springer US, 2003).

46 Nieuwenhuis, B. P. & James, T. Y. The frequency of sex in fungi. Philos Trans R Soc Lond B Biol Sci 371, doi:10.1098/rstb.2015.0540 (2016).

47 Powell, A. E. et al. Fusion between Intestinal epithelial cells and macrophages in a cancer context results in nuclear reprogramming. Cancer research 71, 1497–1505, doi:10.1158/0008-5472.CAN-10-3223 (2011).

48 Lazebnik, Y. The shock of being united and symphiliosis. Another lesson from plants? Cell Cycle 13, 2323–2329, doi:10.4161/cc.29704 (2014).

49 Mani, S. A. et al. The epithelial-mesenchymal transition generates cells with properties of stem cells. Cell 133, 704–715, doi:10.1016/j.cell.2008.03.027 (2008).

50 Qiu, W. et al. No evidence of clonal somatic genetic alterations in cancer-associated fibroblasts from human breast and ovarian carcinomas. Nat Genet 40, 650–655, doi:10.1038/ng.117 (2008).

51 Vitale, I. et al. Multipolar mitosis of tetraploid cells: inhibition by p53 and dependency on Mos. EMBO J 29, 1272–1284, doi:10.1038/emboj.2010.11 (2010).

52 Amend, S. R. et al. Polyploid giant cancer cells: Unrecognized actuators of tumorigenesis, metastasis, and resistance. Prostate 79, 1489–1497, doi:10.1002/pros.23877 (2019).

53 Islam, S. et al. Drug-induced aneuploidy and polyploidy is a mechanism of disease relapse in MYC/BCL2-addicted diffuse large B-cell lymphoma. Oncotarget 9, 35875–35890, doi:10.18632/oncotarget.26251 (2018).

54 Lazova, R. et al. A Melanoma Brain Metastasis with a Donor-Patient Hybrid Genome following Bone Marrow Transplantation: First Evidence for Fusion in Human Cancer. PloS one 8, e66731, doi:10.1371/journal.pone.0066731 (2013).

55 LaBerge, G. S., Duvall, E., Grasmick, Z., Haedicke, K. & Pawelek, J. A Melanoma Lymph Node Metastasis with a Donor-Patient Hybrid Genome following Bone Marrow Transplantation: A Second Case of Leucocyte-Tumor Cell Hybridization in Cancer Metastasis. PloS one 12, e0168581, doi:10.1371/journal.pone.0168581 (2017).

56 Marusyk, A. et al. Spatial Proximity to Fibroblasts Impacts Molecular Features and Therapeutic Sensitivity of Breast Cancer Cells Influencing Clinical Outcomes. Cancer research 76, 6495–6506, doi:10.1158/0008-5472.CAN-16-1457 (2016).

57 Butler, A., Hoffman, P., Smibert, P., Papalexi, E. & Satija, R. Integrating single-cell transcriptomic data across different conditions, technologies, and species. Nat Biotechnol 36, 411–420, doi:10.1038/nbt.4096 (2018).

58 AlJanahi, A. A., Danielsen, M. & Dunbar, C. E. An Introduction to the Analysis of Single-Cell RNA-Sequencing Data. Mol Ther Methods Clin Dev 10, 189–196, doi:10.1016/j.omtm.2018.07.003 (2018).

59 Newman, M. E. Modularity and community structure in networks. Proc Natl Acad Sci U S A 103, 8577–8582, doi:10.1073/pnas.0601602103 (2006).

60 Signorell, A. & al., e. m. (2020).

61 Gatenbee, C. D., Schenck, R. O., Bravo, R. R. & Anderson, A. R. A. EvoFreq: visualization of the Evolutionary Frequencies of sequence and model data. BMC Bioinformatics 20, 710, doi:10.1186/s12859-019-3173-y (2019).

62 Bravo, R. R. et al. Hybrid Automata Library: A flexible platform for hybrid modeling with real-time visualization. PLoS Comput Biol 16, e1007635, doi:10.1371/journal.pcbi.1007635 (2020).

